# Single-Cell Atlas of Early Human Brain Development Highlights Heterogeneity of Human Neuroepithelial Cells and Early Radial Glia

**DOI:** 10.1101/2020.03.06.981423

**Authors:** Ugomma C. Eze, Aparna Bhaduri, Maximilian Haeussler, Tomasz J. Nowakowski, Arnold R. Kriegstein

**Author notes:** These authors contributed equally (Ugomma C. Eze, Aparna Bhaduri).

## Abstract

The human cortex is comprised of diverse cell types that emerge from an initially uniform neuroepithelium that gives rise to radial glia, the neural stem cells of the cortex. To characterize the earliest stages of human brain development, we performed single-cell RNA-sequencing across regions of the developing human brain, including the telencephalon, diencephalon, midbrain, hindbrain, and cerebellum. We identify nine progenitor populations physically proximal to the telencephalon, suggesting more heterogeneity than previously described, including a highly prevalent mesenchymal-like population that disappears once neurogenesis begins. Comparison of human and mouse progenitor populations at corresponding stages identifies two progenitor clusters that are enriched in the early stages of human cortical development. We also find that organoid systems display low fidelity to neuroepithelial and early radial glia cell types, but improve as neurogenesis progresses. Overall, we provide a comprehensive molecular and spatial atlas of early stages of human brain and cortical development.

---

The human brain consists of billions of cells across several functionally interconnected structures that emerge from the neuroectoderm. Many of these structures are substantially expanded or distinct compared to other mammals, particularly the cerebral cortex, the outermost layer of the human brain responsible for perception and cognition. These differences emerge at developmental stages prior to birth, and thus exploring the cell types in the developing human brain is essential to better characterize how cell types across the brain are generated, how they may be impacted during the emergence of neurodevelopmental disorders, and how human neural stem cells can be directed to specific cell types for modeling or treatment purposes.

The brain exponentially increases in size after the neural tube closes^1^. Later in development, across brain regions, a series of similar neurogenic and gliogenic processes give rise to the constituent cell types. However, at the molecular level, the sequence of events that leads to the emergence of these progenitor cells early in development is much less well understood. The diversity of brain structures is known to emerge as a result of segmentation events that generate the prosencephalon, mesencephalon, and rhombencephalon that are then further specified into the anatomical structures (telencephalon, diencephalon etc.) that were dissected in this study^2^ (SFig 1 A-C). It has been proposed that there are core gene regulatory programs that enable the specification of these regions and subsequent development of topographically relevant cell types^3^. We sought to explore if our data could more comprehensively define regional signatures and also identify cell type-specific similarities and differences in these nascent brain structures.

The human cerebral cortex is more than three times expanded compared to our closest non-human primate relatives^4^. The cortex emerges from an initially pseudostratified neuroepithelium that gives rise to radial glia, the neural stem cells of the cortex^1^. Radial glia generate neurons, initially through direct neurogenesis, and then indirectly through transit-amplifying intermediate progenitor cells^5^. A number of subtypes of radial glia have been identified as the cortex matures, and their primary role in neurogenesis declines late in the second trimester, at which point they generate the glial populations of the cortex^6^. scRNA-seq has added substantially to our knowledge about cellular diversity and signaling networks, particularly during stages of peak neurogenesis. However, the first trimester of cortical development has not been described at this level of molecular detail, and important questions remain about the timing of neurogenesis, the presumed uniformity of the neuroepithelium, and the signals that promote the transition to radial glia.

## Results

### Whole Brain Analysis

In order to identify cell types and trajectories that lay the foundation for the development of the human brain, we performed single-cell RNA sequencing (scRNA-seq) using the droplet-based 10X Genomics Chromium platform. We sequenced cells from ten individuals during the first trimester of human development, spanning Carnegie Stages (CS) 12 to 22, corresponding to gestational weeks (GW) 6 – 10. We also included cortical samples from one CS13 and one CS22 individual that were analyzed in a prior study^7^. In order to identify the major cell populations across brain regions and to compare them to one another, we sampled all available and identifiable structures, including the telencephalon, diencephalon, midbrain, hindbrain, cerebellum, ganglionic eminences, thalamus, hypothalamus, and cortex (SFig 1 – 3, STable 1). We validated that our sequencing did not contain substantial artefacts or cell debri, and that it represented highly expressed transcripts from bulk RNA-sequencing experiments^8^ (SFig 1). In total, we collected 289,000 cells passing quality control.

Across brain regions, hierarchical analysis initially identified two major cell classes – progenitors and neurons. To identify the developmental region-specific gene signatures for each brain area, including the hindbrain, midbrain, thalamus, ganglionic eminences, and cortex, we performed differential gene expression across areas at each age (STable 2 – 4). Using samples at CS22, we generated data-driven regional signatures and explored when the most structure-specific genes emerged. We observed that characteristic transcription factor expression (such as *HOX* genes [hindbrain]^9^; *PAX7* [midbrain]^10^; *GBX2* [thalamus]^11^; *NKX2-1* [medial ganglionic eminence]^12^; and *FOXG1* [cerebral cortex]^13^) segregated these regions from one another as early as CS13. However, the cell type-defining gene expression programs were largely conserved across brain regions, resulting in subsets of progenitors or neurons that transcriptomically appear similar across regions (SFig 2B). This was reflected by the fact that co-clustering at the early stages could not segregate regional identities, but that this became possible at later stages. At the earliest timepoint, CS12, the differences between brain regions were minimal and did not resemble the more advanced region-specific programs that were identifiable later in the first trimester (SFig 2 – 3).

### Single-cell Sequencing of the Telencephalon

To better understand the early stages of cortical development, we focused on the 59,000 cells in our dataset that originated from the telencephalon (earliest dissections) and the cortex specifically (when it was identifiable for subdissection). Clustering this data revealed minimal batch effects as most clusters contained contributions from multiple individuals, and clustering segregated the samples based on early and late first trimester stages (Figure 1, SFig 4). Each of the 63 identified clusters could be assigned to a cell type identity of either neuroepithelial cells, radial glia, intermediate progenitor cells (IPCs), neurons, or mesenchymal-like cells. To differentiate between the radial glia and neuroepithelial cells in our transcriptomic data, we examined if *SOX2*-positive progenitor clusters contained evidence of neurogenic genes, considered to be a criteria of radial glia identity^14^. *SOX2*-positive progenitors that did not have these characteristics were labeled neuroepithelial cells. Additionally, we noted “other” cell populations of endothelial cells, microglia, and pericytes (STable 5 – 6). Furthermore, we observed that the mesenchymal cell population diminished strikingly over the course of the first trimester. Comparison of these clusters to previously published single-cell data, including a small number of first trimester cells, showed a strong cluster-level correlation, suggesting that our larger dataset presented here recapitulated populations that were previously observed. Additionally, in our previous analysis, some clusters from these early samples were marked “unknown” in identity^15^; our correlation analysis is sufficient to now assign cell type identities to these clusters (SFig 5).

**Figure 1.**
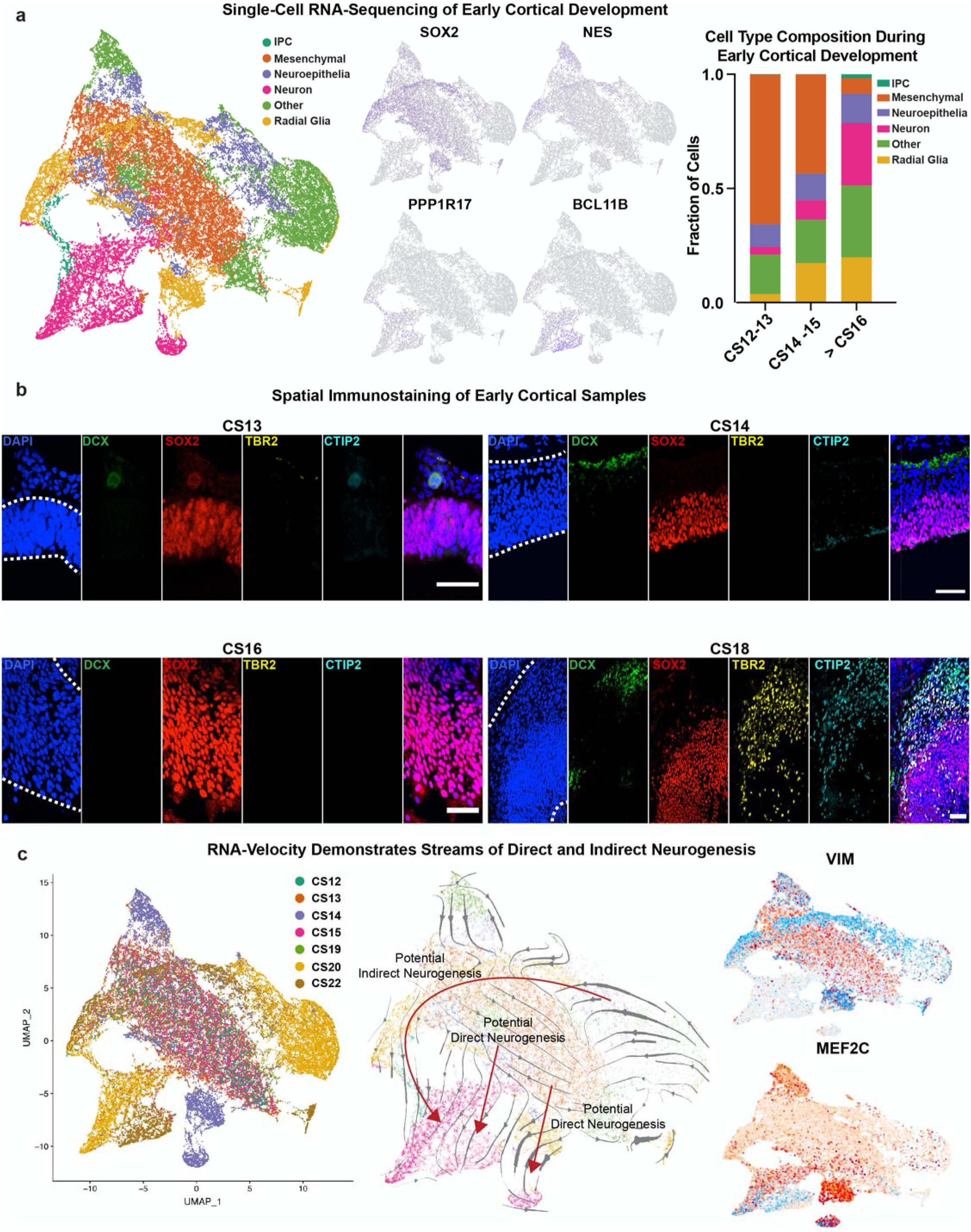
Cell Types in the Early Human Cortex. **a)** UMAP plot of 58,145 telencephalon or cortical dissections colored by annotated cell type. Feature plots of markers of broad progenitors (*SOX2*), radial glia (*NES*), IPCs (*PPP1R17*), and neurons (*BCL11B*) are shown. Stacked bar chart shows cell type composition at earliest (CS12-13), middle (CS14-16), and late (CS19-22) first trimester. **b)** Immunostaining for major cell type markers across first trimester stages. Because of limited sample availability, each sample was immunostained one time. Nuclei shown in blue (DAPI, dotted line demarcates cortical span), newborn neurons marked by doublecortin (DCX, green), progenitors by SOX2 (red), IPCs by TBR2 (yellow), and maturing neurons by CTIP2 (cyan). Scale bars are 50 μM, except CS13 which is 25 μM. **c)** UMAP colored by age shows the segregation of samples by early and late first trimester stages. RNA-velocity trajectories are depicted by gray arrows in the middle UMAP plot with underlying color by cell types as annotated in (a). Thickness of the lined arrow indicates the differences in gene signature between cell types, and red arrows show predicted direct and indirect neurogenesis trajectories. The velocity plots on the right show the intensity of scored velocity for a progenitor gene, *VIM*, and a neuronal gene, *MEF2C*. Velocities highlight the distinction between progenitors and neurons in the clustering and velocity analyses.

We observed that although neurogenesis ramps up by the end of the second trimester, even the earliest samples contained a small number of neurons (Fig 1A). It seemed unlikely that they could be migratory populations from other cortical regions since migratory interneurons have not been identified until later developmental timepoints^16^. Presumably these neurons were produced locally by direct neurogenesis from radial glia which occurs early in non-human models of cortical development as well as human cortical organoids^17^, and would be consistent with the observation that the IPCs that mediate indirect neurogenesis are nearly absent until later in the first trimester (after CS16). However, without lineage tracing, the inference that the earliest neuronal populations are a product of direct neurogenesis remains a working hypothesis. Sub-clustering of the neuronal populations resulted in subtype clusters, some of which were strongly enriched in either younger samples (< CS16, presumed to result from direct neurogenesis) or older samples (> CS16) (SFig 6, STable 7). These included expected differences such as clusters marked by *NHLH1*, which was significantly higher in the older samples, and has been associated with newborn neurons derived from IPCs^18^. Other genes significantly enriched in older samples included *NEUROD6* and *BCL11B* which are associated with neuronal and deep layer identity^14^, as well as *CALB2* which is expressed in migratory ventrally-derived interneurons and a subset of excitatory neurons^19^. We noted significantly higher expression of *MEF2C* in the younger samples. This has been identified as a regulator of early neuronal differentiation and layer formation^20^, but is also a factor involved in synaptic maturation at later stages^21^. We validated high expression of *MEF2C* at the earliest timepoints using *in situ* hybridization, though some of the expression was extra-cortical. By CS22 its expression had diminished, but *MEF2C* was highly expressed again by mid-gestation at GW14 (SFig 7). This expression pattern is intriguing given the role of MEF2 transcription factors in regulating apoptosis^22^.

We also observed clusters marked by previously undescribed genes, including the younger sample-specific *LHX5-AS1* cluster. We validated that *LHX5-AS1* was strongly enriched at these early time points with broad expression at CS13, but restriction to the developing cortical plate by CS14 and CS16 (SFig 8).

Interestingly, *LHX5-AS1* RNA had the same expression pattern as LHX5 protein (SFig 9), indicating that it may play a repressive role to the protein, which has been characterized in Cajal-Retzius cell development^23^ but has largely remained unstudied in these early cell populations. The remaining clusters consisted of expected early-born neuronal populations including Cajal-Retzius cells marked by *RELN* (SFig 10), and subplate cells marked by *TLE4* and *NR4A2*. The subplate clusters were unexpectedly heterogeneous (SFig 6C-D) and contained marker genes not previously associated with subplate identity (STable 7), suggesting our analysis may provide additional characterization of early neuronal cell type gene expression patterns.

We also sought to describe the spatial organization of the cortical populations across development. Thus, we performed immunostaining validation in five individuals from the first trimester compared to an early second trimester sample at GW14. We observed FOXG1 staining as early as CS16, but not earlier, as has been described previously^24^ (SFig 11). FOXG1 staining ensured that we were identifying cell types in the developing forebrain. We further defined cortical regional and progenitor identity by positive PAX6 expression and positive SOX2 expression. PAX6 expression has previously been identified as a determinant for neuroectoderm fate^25^ and SOX2 as a marker for stem cells. We were also confident of cortical identity based on the anatomical presence of the optic cups (SFig 12) just ventral, lateral, and caudal, as well as the nasal ridge, (SFig 12) located just ventral to our regions of interest. Tilescan images of these markers are available for download and exploration in the image browser we created in conjunction with this study [https://cells-test.gi.ucsc.edu/?ds=early-brain, Images tab].

In concordance with our transcriptomic data, we observed prevalent SOX2 staining as early as CS13, that persisted through GW14. Newborn neurons marked by DCX were identifiable as early as CS14, but markers of maturing neuronal identity such as BCL11B (CTIP2) emerged after CS16. As expected, IPCs identified by TBR2 staining did not emerge until CS18 in the dorsal telencephalon^26^ (Fig 1B, SFig 13), though we and others have observed TBR2 staining at earlier timepoints (CS16) in the ventral telencephalon^27^.

### RNA-Velocity Analysis of Lineage

In order to characterize the lineage relationship between major cortical cell types, we performed RNA-velocity analysis using the scVelo algorithm^28^. This algorithm incorporates mRNA levels and inherent expression variability of individual cells to infer steady states that contribute to potential lineage relationships. Across cortical cell types, the most apparent lineage stream originated in progenitor cell populations from older samples and followed a trajectory through IPCs to neurons (Fig 1C). This trajectory of neurogenesis is well described and suggests that the stereotypical process of neuronal differentiation emerges after CS19. At earlier time points, there were local examples of radial glia giving rise to neurons, suggestive of potential direct neurogenesis, and also lineage relationships between the progenitors themselves. In addition to velocity streams related to the excitatory lineage, there was also a stream from a collection of putative blood-brain barrier cells toward the middle mesenchymal mass. Given the age of these samples, this stream is suggestive of the onset of vasculogenesis which occurs at this time^29^. This is supported by further velocity analysis demonstrating an increase in velocity (indicated in red) of presumptive endothelial (*FN1*) and pericyte (*RGS5*) markers, but not markers of microglia identity (*AIF1*) or the marker of the mesenchymal population (*LUM*) (SFig 4D).

The transition from neuroepithelial cells to radial glia has traditionally been characterized by the reorganization of tight junctions and the appearance of nestin RC2 (NES) immunoreactivity^30^. We sought to visualize this process and its transition across our samples. Using NES as a marker of radial glia, TJP1 (ZO-1) as an indicator of neuroepithelial cells, and SOX2 as a label for all progenitor populations, we saw small numbers of nestin-positive radial glia at CS14, with substantial upregulation by CS22 that simultaneously corresponded to a waning of ZO-1 (Fig 2A). By GW14, ZO-1 staining dissipated and was largely constrained to presumed vascular structures. During the first trimester we found a progressive, and incremental, shift away from neuroepithelial identity (SFig 14). In order to better understand the heterogeneity and spatiotemporal trajectories of neuroepithelial and radial glia progenitor populations, we sub-clustered the progenitors. Because the mesenchymal population also expressed strong progenitor cell markers including *VIM* and *SOX2*, we removed all neurons, IPCs, and other cell populations (including microglia, pericytes, and endothelial cells) and sub-clustered the remaining cells. The resulting analysis yielded nine sub-clusters all marked by strong *VIM* and *SOX2* expression (Fig 2B). Each of the nine clusters contained some cells from all first trimester age samples, suggesting that the velocity and maturation trajectories were not purely a function of age (SFig 15B, STable 8). To examine if these trajectories correspond to expected cell identity transformations from neuroepithelial cells to radial glia, we explored the expression of *HES5* and *FGF10*, which have been described to mediate the transition between these progenitor populations^31,32^. As expected, *HES5* was strongly enriched in radial glia compared to other cell types, and *FGF10* peaked in neuroepithelial populations (SFig 15). We explored if any of these progenitors expressed gene signatures that defined excitatory neurons across cortical areas, and found that three-quarters of the progenitors uniquely expressed an areal identity while the rest were unspecified (SFig 15). We additionally performed WGCNA analysis of the progenitor clusters as an orthogonal metric to identify distinct populations^15,33^. We found strong correspondence between clusters derived from WGCNA analysis and our nine progenitor subtypes, and also found strong enrichment for radial glia-like networks in later stage samples (SFig 15, STable 12).

**Figure 2.**
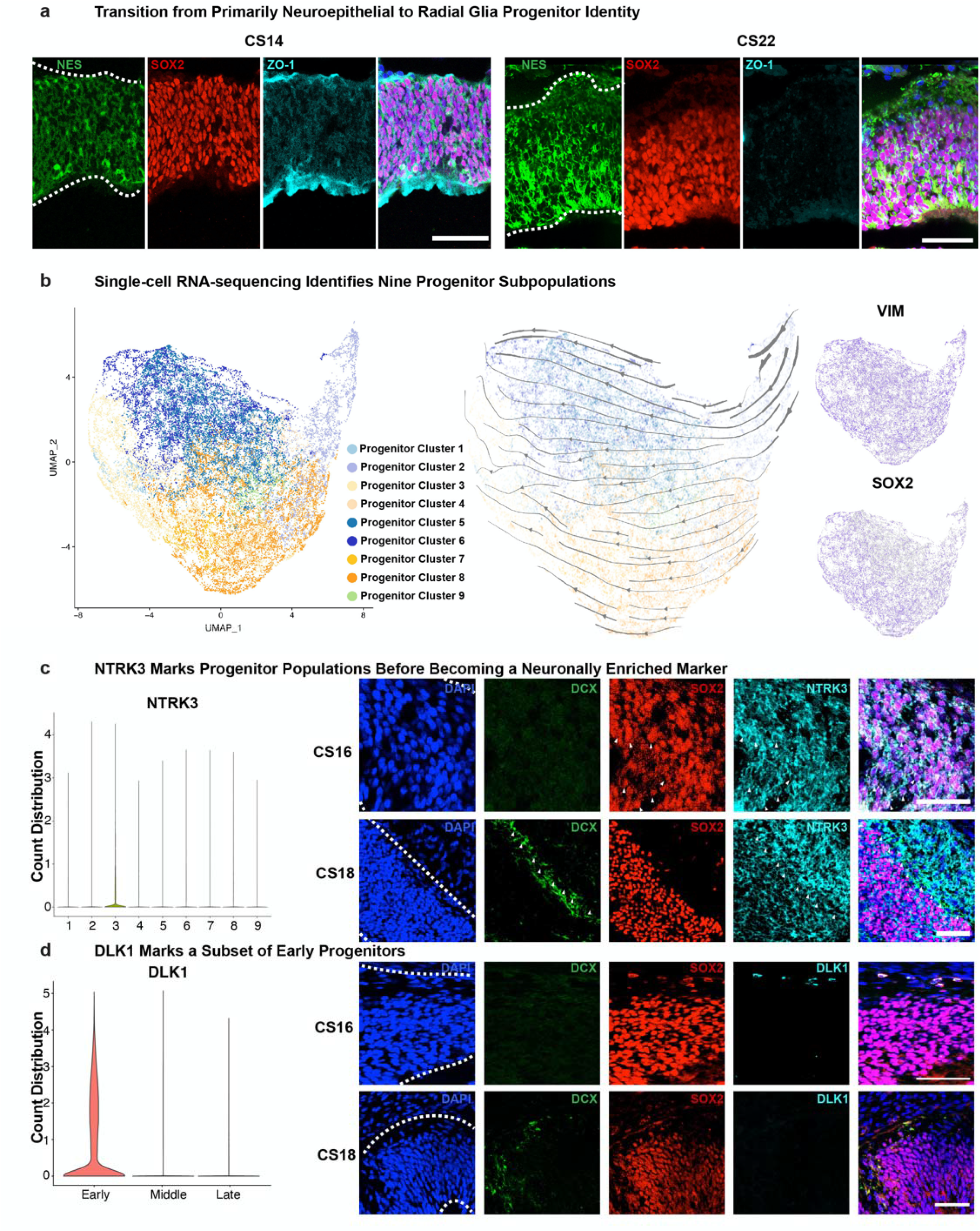
Early Progenitors Can be Divided into Nine Progenitor Subtypes. **a)** Immunostaining of early first trimester samples (including CS14, shown here) show strong staining for all progenitors (SOX2, red), as well as tight junctions (ZO-1, cyan), but limited staining for radial glia (nestin, NES, green). By CS22, NES staining increases substantially, ZO-1 decreases, and SOX2 expression is maintained. Scale bars are 50 μM. **b)** Left UMAP plot depicts subclustered progenitor cells, while the middle plot shows the velocity trajectory across progenitor subtypes. Feature plots on the right show high expression of *VIM* and *SOX2* marking all progenitor populations. **c)** Progenitor Cluster 3 is specifically and uniquely labeled by *NTRK3* as shown in the violin plot. Immunostaining for NTRK3 (cyan) shows early labeling of SOX2 (red) positive progenitors at CS16 (white arrows), but expression shifts to more closely coincide with newborn neurons marked by DCX (green) by CS18 (white arrows). Scale bars are 50 μM. **d)** DLK1 (cyan) is the top marker for Progenitor Cluster 8, and is exclusively expressed in early first trimester as shown in the violin plot on the left. Immunostaining for DLK1 at CS16 shows co-localization with low SOX2 (red) expressing cells at the boundary of the cortical edge. This staining disappears from the cortex entirely by CS18 when DCX (green) staining emerges. For all panels, each sample was immunostained one time.

To examine if this progenitor distribution was distinct at the earliest timepoints of our dataset, we sub-clustered the CS12 and CS13 samples individually. From this analysis we identified 29 populations that were characterized by *SOX2* (progenitor), *TOP2A* (dividing), *LHX5-AS1* (previously uncharacterized cell population), and *LUM* (mesenchymal cells). Comparison of these subpopulations to the nine progenitor subtypes identified across all samples demonstrated strong correspondence to these nine cell types, with additional heterogeneity in the CS12 and CS13 mesenchymal populations (SFig 5B, STable 9). Velocity analysis identified a strong gradient across the populations from cluster 2 towards clusters 1 and 6. These data strongly suggested a gradient of identity across the nine progenitor populations (SFig 15D). We characterized the genes across all our velocity analyses that most strongly influenced the velocity stream (and thus would be hypothesized to be the chief regulators of cell fate transition). The drivers of velocity were enriched for genes that have been associated with neurodevelopmental or psychiatric disorders, including autism, cortical malformations, and schizophrenia (STable 10), indicating that very early events in cortical development may result in susceptibility or vulnerability to these disorders. More work at these early timepoints is required to fully characterize the developmental implications.

### Progenitor Heterogeneity

In order to explore the timing and spatial localization of each of the progenitor subpopulations, we performed immunostaining of the most specific gene markers for each cluster (SFigs 7 – 10; SFigs 16 – 26). Each of these immunostainings is available as a tilescan of the entire brain section in our image browser [https://cells-test.gi.ucsc.edu/?ds=early-brain, Images tab]. *NTRK3* was a highly specific marker of Progenitor Cluster 3, and intrigued us because it is commonly described as a TrkC receptor that enables survival of specific neuronal populations^34^ and has broad neocortical expression at later stages of mouse cortex development^35^. We observed that at early stages (CS14, CS16), NTRK3 broadly co-localized with SOX2-positive progenitor cell populations (Fig 2C, SFig 16), but after CS18, it shifted from progenitor to neuronal expression as would be expected. *DLK1* was a top marker gene for Progenitor Cluster 8 and also was highly enriched in early samples. Though much of the DLK1 staining was in stromal cells peripheral to the cortex, as has been previously reported^36^, we observed a small number of DLK1-positive cells that expressed low SOX2-levels at early stages, such as CS16, after which DLK1 expression completely disappeared (Fig 2D, SFig 17). DLK1 is a non-canonical ligand of Notch that inhibits its function and impacts Notch 1 receptor distribution^37^, though it is not widely expressed in the mammalian central nervous system during development^38^. DLK1 has also been described to play a role in enabling postnatal neurogenesis in the mouse^39^, and may play a similar regulatory role at the earliest neurogenic timepoints of human cortical development. Both NTRK3 and DLK1 may play important roles in receptor/ligand communication, as integration with other brain regions increases (STable 11). Across progenitor populations we describe known and new gene expression patterns that define the neuroepithelial and radial glia subpopulations and suggest that there exists an expression gradient that marks the transition from early to more mature progenitor populations.

Both in our initial clustering and in the more focused progenitor analysis, a mesenchymal-like population labelled by the gene *LUM* was distinctly segregated from other progenitor populations. Lumican, the protein encoded by *LUM*, is an extracellular matrix protein that is widely present in mesenchymal tissues throughout the adult body^40^. It has also been used to promote folding in cortical structures when added exogenously to media^41^. In our data, *LUM* expression is highly enriched in the mesenchymal cell population and diminishes substantially after early developmental timepoints (Fig 3A). Another marker of this cluster was *ALX1*, which also has one of the most correlated gene expression patterns to *LUM* (Fig 3B). ALX1 has been described as required in knockout mice for the development of the forebrain mesenchyme^42^, but has not been studied in humans. We observed prevalent LUM staining in our samples at or prior to CS16, with ALX1 at the edges of the forebrain, including several co-localized PAX6-positive cells (Fig 3 C-D, SFig 18 – 19). ALX1 staining significantly diminishes later in development.

**Figure 3.**
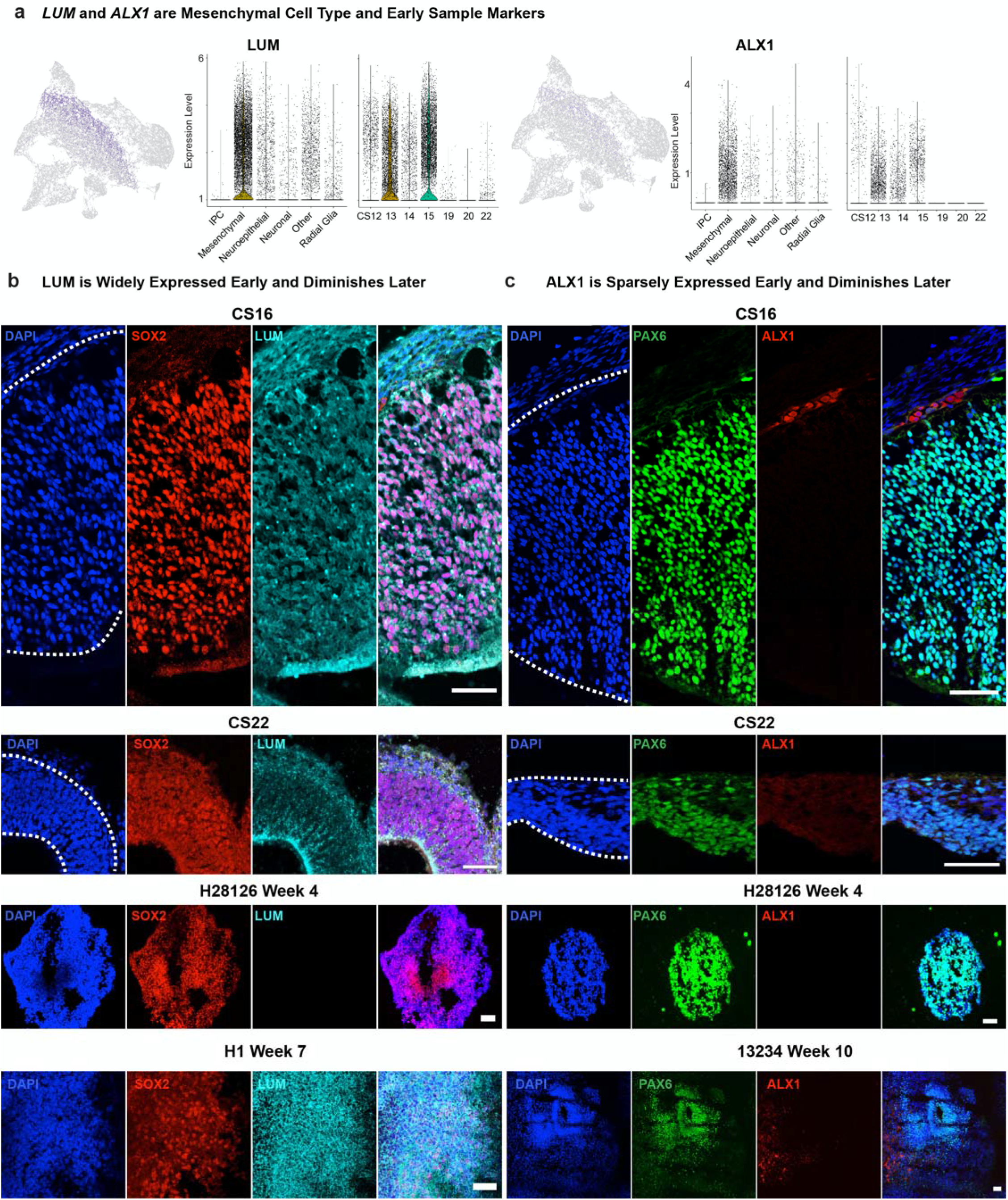
Single-cell RNA-sequencing Identifies Early Mesenchymal Cell Population. **a)** In both progenitor and cortex clusterings, *LUM* marks a separate population of cells, as shown by the feature plot on the l ft. *LUM* expression is highly specific to the mesenchymal cell type and is enriched in early samples. *ALX1* expression is highly correlated to *LUM* as shown in the right feature plot, and is similarly enriched in early, mesenchymal populations. **b)** Immunostaining for LUM (cyan) shows prevalent expression in and between progenitors marked by SOX2 (red) at CS16, but this expression dissipates by CS22. However, expression of LUM does not begin until Week 7 in the H1 organoid. Scale bars are 50 μM. **c)** Immunostaining for ALX1 (cyan) shows sparse expression in PAX6 (green) positive cells, that disappears from the cortex but is expressed in surrounding brain structures at CS22. ALX1 expression does not begin until Week 10 in the 13234 cerebral organoids. For all panels, each sample was immunostained one time. Scale bars are 50 μM.

### Canonical Developmental Signaling Pathway Activation

Because canonical signaling pathways have been characterized for their role in patterning the human brain and cortex, we sought to explore the dynamics of their expression patterns in human cortical progenitors. scRNA-seq data showed dynamic expression of the genes that have been described as part of the FGF, Wnt, mTOR and Notch signaling pathways (Fig 4A) in cortical progenitors. The FGF signaling family has been described to assign rostral identity in the developing neural tube. But FGF also plays a role in the transition to and maintenance of the radial glia progenitor pool, in part promoted by FGF10, by permitting the transition of neuroepithelial cells to radial glia and by preventing the transition into intermediate progenitors^32^. Furthermore, FGF signaling interacts with the Notch signaling pathway during cortical development. Classic studies have demonstrated that Notch signaling is generally required to preserve stem cell pools. Constitutive Notch 1 activation increases radial glial generation and leads to a decrease in the expression of pro-neural genes, like Neurogenin-2 (Ngn2)^43^. Consistent with this observation, here we find that Notch 1 staining peaks and is largely restricted to the ventricular zone at CS14 and CS16 (Fig 4D).

**Figure 4.**
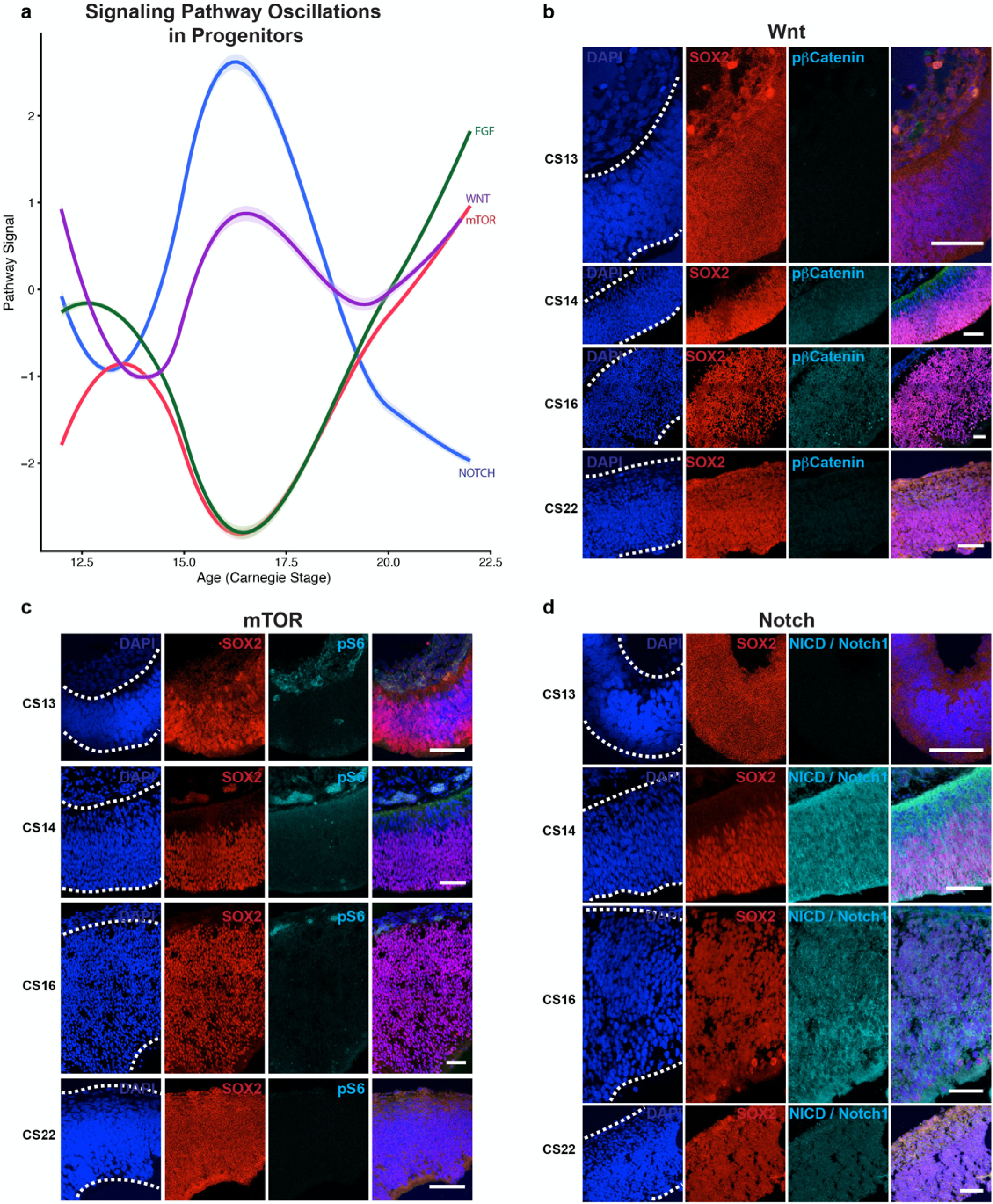
Signaling Pathway Oscillations in the First Trimester Human Cortical Progenitors. **a)** RNA expression patterns of the FGF (green), Wnt (purple), mTOR (red) and Notch (blue) signaling pathways as defined by KEGG pathway designations in progenitors across ages sampled in this study. The light same colored shading surrounding each bar indicates the loess regression 5-95% confidence interval. **b)** Wnt activity, as indicated by phosphorylated β-catenin (cyan), is highest in the CS14 and CS16 samples and colocalizes with progenitors (SOX2, red), but dissipates by CS22. **c)** mTOR activity, as indicated by phosphorylated S6 ribosomal protein (cyan), primarily localizes along the cortical plate (DCX, green) in the youngest samples and diminishes by CS22. **d)** Notch activity, as indicated by cleaved Notch intracellular domain (NICD) of Notch 1, peaks in expression at CS14 and localizes mainly to the ventricular zone. Nuclei are labeled by DAPI in blue. For all panels, each sample was immunostained one time. All scale bars are 50 μm.

mTOR activity in the developing forebrain has largely been uncharacterized at early developmental timepoints, and descriptions of its function have been restricted to its role in regulating oRG cells^15^. However, upregulation of CDC42-dependent mTOR signaling has been shown to be sufficient to generate neural progenitors, mediated through increased HES5 and PAX6 expression^44^, suggesting that mTOR signaling early on may be important for driving the switch from neuroepithelial stem cell to radial glia. We find that expression of phosphorylated S6 is highest in the earliest samples and then diminishes later in the first trimester (Fig 4C), further supporting a role in the neuroepithelial to RG fate transition. Furthermore, Wnt signaling has also been implicated in medial-lateral patterning of the neural tube and in cell fate transitions. Constitutive Wnt activation leads to opposing actions of increasing the progenitor pool by preventing neurogenesis but also of increasing the neuronal pool by promoting IPC differentiation^45^. This suggests that Wnt activity is variable and cell-dependent in cortical development. In this study we discover that active Wnt signaling (phosphorylated β-catenin) peaks at CS16 and diminishes subsequently (Fig 4B). These data further clarify our understanding of signaling in patterning cortical progenitors, with important implications for better modeling early stages of cortical development.

### Conservation of Progenitor Populations Across Species

In order to evaluate the similarities and differences between early forebrain development in human and rodent, we performed scRNA-seq of the mouse forebrain at embryonic days (E) 9 and 10, which correspond to Theiler Stages (TS) 14 and 16. Although the onset of neurogenesis differs between human and mouse^46^, we used immunostaining to verify that at these mouse stages, the forebrain was already expressing FOXG1 and undergoing neurogenesis, as marked by the presence of DCX (SFig 11, SFig 13). To compare mouse and human progenitor populations, we performed clustering and correlation analysis between the mouse clusters and our nine human progenitor populations. Using a correlative threshold, we observed that seven of the nine human populations had at least one corresponding cluster in the mouse single-cell data (Fig 5A). However, there were no high correlations for Progenitor Cluster 4 (marked most highly by *C1orf61*) or Progenitor Cluster 7 (marked most highly by *ID4*). To test the hypothesis that these populations were underrepresented in the mouse data, we used immunostaining to explore the expression of the major progenitor population markers in the mouse. Indeed, no fluorescent *in situ* signal for *C1orf61* or immunostaining signal for Id4 was identifiable in the mouse forebrain at TS14 or 16 (Fig 5B, SFig 20 – 22). Although the TS14 and 16 samples were more advanced in terms of neurogenesis, they may be at timepoints that are relatively immature in other ways compared to our first trimester samples. To account for this difference, we also explored the single-cell trajectory of *Id4* from E9 – E10 (our data) to E11.5 – E17.5 (Yuzwa et al, 2017)^47^. In the mouse, *Id4* peaks at E11.5 and is low after E13, whereas human *ID4* is sustained in its expression pattern after CS13 (Figure 5D). By contrast, both *C1orf61* RNA and ID4 protein were detectable in macaque embryonic tissue and chimpanzee induced pluripotent stem cell derived organoids (SFig 23). C1orf61 (CROC-4) has been characterized to be widely expressed during late developmental stages and in the adult brain, regulating c-FOS signaling^48^, but is otherwise uncharacterized. In contrast, Id4 has been identified in the developing mouse cortex at later stages (E15.5). ID4-deficient mice exhibit impaired brain size and mistimed neurogenesis^49^, suggesting that earlier expression in human progenitors compared to mouse could indicate a potential mechanism by which early progenitors expand more rapidly in humans than in mice.

**Figure 5.**
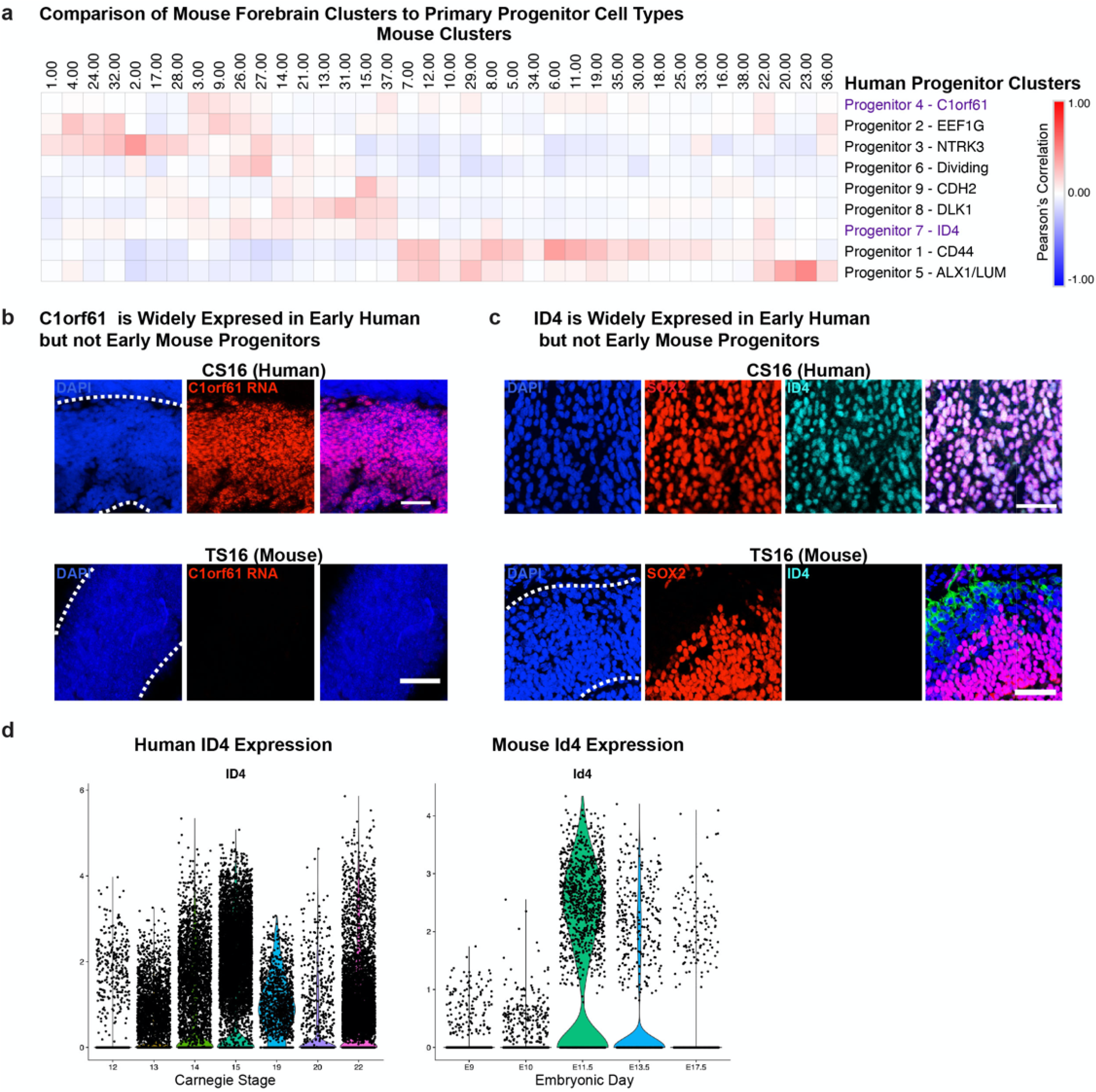
Mouse Models Different Aspects of Early Cell Types. **a)** Heatmap showing the comparison of mouse forebrain clusters from 16,053 cells to the primary progenitor populations identified in this dataset. Correlations are performed using Pearson correlations between cluster marker sets and identify Progenitor Clusters 4 and 7 to not have a counterpart in mouse data. **b)** Fluorescent *in situ* hybridization of human samples at CS16 shows broad expression of C1orf61 (cyan) in progenitor cells labeled by SOX2 (red). Parallel *in situ* staining in TS16 mouse shows no C1orf61 expression. Scale bars are 50 μM. **c)** Immunostaining of human samples at CS16 shows broad expression of ID4 (cyan) in progenitor cells labeled by SOX2 (red). A parallel staining in TS16 mouse shows no ID4 expression. Scale bars are 50 μM. **d)** Violin plots of ID4 RNA expression across several Carnegie stages (human) and embryonic days (mouse). ID4 RNA expression in the human persists onward from CS13. However, ID4 RNA expression in the mouse peaks at E11.5 (TS20) and dissipates. For all panels, each sample was immunostained one time.

Despite the general conservation of cell types between the early development of the human and mouse cortex, the absence of specific human cell populations at early developmental stages in the mouse suggests that alternatives to mouse are required to fully model the earliest cell types involved in human cortical development. Cortical organoids are an attractive model because they can be generated from normal or patient-derived induced pluripotent stem cells (iPSCs) and are amenable to genetic modulation. We previously generated an extensive catalog of single-cell sequencing data from multiple organoids derived from four pluripotent stem cell lines using three protocols analyzed from week 3 to 24^7^. Using 108,593 cells derived from week 3 and 5 of this dataset, we explored the fidelity of organoid cell types to early human cortex. The preservation of excitatory neuronal identity mirrored that of later developmental stages (∼0.5, as we recently reported). However, the remaining primary cell types had much lower fidelity in organoids (average of 0.28) (Fig 5C). These data suggest that early cell types in organoids do not resemble their corresponding cortical progenitor counterparts, although the fidelity of cell types does improve at later stages of organoid culture. Interestingly, the two populations that were not well represented in mice at TS14 and 16 were identifiable in organoid cultures as evidenced by fluorescent *in situ* probing for C1orf61 and immunostaining for ID4 (SFig 23).

## Discussion

Here, we present a comprehensive overview of single-cell RNA-sequencing from the first trimester of human development. As these samples are rarely accessible, we seek to present both the single-cell sequencing as well as the immunostaining validation provided as part of this study as a community resource. Our analysis highlights the granular gene expression programs that emerge across brain structures and across cell types in the human neocortex as development progresses, and provides a dataset by which other model organisms and *in vitro* cortical organoids can be compared to their primary human counterparts. Importantly, the tissues used for the purpose of this study are fragile, and dissections rely on morphological hallmarks without knowledge of expression patterns prior to sample collection. As such, the single cell analysis becomes even more essential as a tool to explore and validate accurate regional identities. The gene signatures that we describe and validate may offer additional markers that can be used to mark these brain regions, as well as the transition from neuroepithelia to radial glia cell identity.

Our analysis of several brain regions highlights many similarities across progenitors that were only mildly distinguished by the expression of regional-specific transcription factors. These similarities in gene programs across the developing human brain suggest that the process by which initially uniform stem cells give rise to neuronal and glia heterogeneity characteristic of the adult brain are parallel across brain structures and that insights from one region, including the degree of heterogeneity and the trajectories of differentiation, may be able to cross-inform identification of regulatory gene programs across the brain. We aspired to annotate specific gene programs that distinguish progenitor groups within the cerebral cortex. Instead, we observed a gradient of expression patterns, with clear neuroepithelial and radial glia cell populations on the ends of the spectrum, indicating that the transition from one population to another may be gradual rather than distinct. However, because our data is limited by the snapshot view of each time point sampled, and by imperfect representation of all possible timepoints along this continuum, this conclusion will need to be further explored through additional sampling, validation, and eventual mechanistic examination.

One unexpected population that was identified in our progenitor dataset was a population that appeared mesenchymal in origin (marked by ALX1 and LUM) by both single-cell sequencing and immunostaining, that comprised a large swath of the telencephalon at the earliest timepoints that we sampled. We hypothesize that the ALX1-positive cells may resemble previously described neural crest-derived cell populations that give rise to the meninges^50^. These cells may be secreting LUM as a structural component to support the physical formation of the telencephalon prior to the emergence of the radial glial scaffold. In support of this hypothesis, we found that the ALX1-positive cells were in the presumptive stroma surrounding the cortex, while LUM was detected throughout the cortex. Although cortical organoids do not express ALX1 or LUM at early stages, their expression emerges after seven weeks of development (Fig 3 B-C), further prompting questions about their function and role in cortical development.

Comparisons of primary data presented here to the single-cell sequencing of previous cortical organoid populations raises interesting differences between the systems. We were surprised to find limited cell type correspondence at the earliest stages of cortical organoid generation, and increased fidelity once the radial glia and neuronal populations emerged. This may indicate that terminal identity does not depend upon a specific differentiation path. However, there were also differences in the timing and cell type composition between early human and mouse populations, further highlighting that no model system is ideal for the study of all biological questions. Moreover, populations present in primary human tissue that were not present in early organoids, including the *LUM* and *ALX1* clusters, could be identified in mice (SFig 18 – 19), indicating that depending on the questions being investigated, either *in vivo* mouse or *in vitro* human models may be most appropriate for the study of neuroepithelial and early radial glia populations. Our description of early human cortical cell populations may also enable further refinement of *in vitro* culture systems to better reflect human neuroepithelial and radial glia populations. Together, the data we present here represent a characterization of major cell types across the first trimester of human brain development and highlight the subpopulations of progenitor cells that form the basis for creating the human cortex.

### Accession Codes

The data analyzed in this study were produced through the Brain Initiative Cell Census Network (BICCN: RRID:SCR_015820) and deposited in the NeMO Archive (RRID:SCR_002001): https://assets.nemoarchive.org/dat-0rsydy7.

## Acknowledgments

The authors thank S. Wang, W. Walantus, M.G. Andrews, G. Wilkins, L. Subramanian, A. Pollen, M. Speir, and members of the A.R.K. laboratory for providing resources, technical help and helpful discussions. Single-cell RNA sequencing data has been deposited at the NeMO archive under dbGAP restricted access. All primary human tissue was obtained from the Human Developmental Biology Resource (HDBR), with special thanks to S. Lisgo and M. Crosier. This study was supported by NIH award U01MH114825 to A.R.K., and F32NS103266, K99NS111731, and L’Oreal For Women in Science Award through the AAAS to A.B. A.B. and A.R.K. designed the study and analysis. Experiments were performed by U.C.E., A.B., and T.J.N. Data analysis was performed by U.C.E., and A.B. The study was supervised by A.B. and A.R.K. This manuscript was prepared by A.B. and U.C.E. with input from all authors. A.R.K. is a co-founder and Board member of Neurona Therapeutics. The remaining authors have no competing financial or non-financial interests.

## Online Methods

Additional methodological details can also be found in the Life Sciences Reporting Summary included in the Supplementary Material.

### Sample Processing

Acquisition of all primary human tissue samples was approved by the UCSF Human Gamete, Embryo and Stem Cell Research Committee (GESCR, approval 10-03379 and 10-05113). All experiments were performed in accordance with protocol guidelines. Informed consent was obtained before sample collection and use for this study. First-trimester human samples were collected from elective pregnancy terminations through the Human Developmental Biology Resource (HDBR), staged using crown-rump length (CRL) and shipped overnight on ice in Rosewell Park Memorial Institute (RPMI) media or in 4% paraformaldehyde. Samples were donated and deidentified for sex. Although we can infer sex for sequenced samples, we have not accounted for it because it is an unclear measurement during the first trimester. Fixed samples were used for downstream immunostaining and imaging. Mouse samples (CD-1® IGS Mouse) were sacrificed at E9 and E10, and staged using the somite number. Mice were housed in shared housing, 5 mice to a cage with a 12 light/12 dark cycle and Temperatures of 65-75°F (∼18-23°C) with 40-60% humidity. Equal numbers of male and female embryonic mice were used. All mouse experiments were approved by and conducted according to the UCSF Institutional Animal Care and Use Committee (protocol AN078703-03A). Half of the mouse samples were randomly assigned for fixation in 4% paraformaldehyde and the remaining live samples were used for dissociation. Live samples were subdissected into identifiable regions and dissociated using papain (Worthington, LK003150) with DNase. Samples were minced and incubated in 1ml activated papain for 15 min at 37°C, according to the manufacturer’s instructions. Samples were then inverted for several times and incubated for an additional 15 min. The dissociated cells were centrifuged at 300 x *g* for 5 min and the papain was removed. Macaque tissue sections (E64) were a gift from A. Pollen (UCSF), originally a gift from A. Tarantal (UC Davis).

### Single-cell RNA Sequencing

Single-cell capture was performed following the 10X v2 Chromium manufacturer’s instructions. Each sample was its own batch. For each batch, 10,000 cells were targeted for capture and 12 cycles of amplification for each of the cDNA and library amplifications were performed. Libraries were sequenced according to the manufacturer’s instructions on the Illumnia NovaSeq 6000 S2 flow cell (RRID:SCR_016387).

### Single-cell RNA Analysis

Single-cell RNA-sequencing data was aligned to the GRCh38-0.1.2 reference genome, and cells were identified using CellRanger v2 (RRID:SCR_017344). Quality control removed cells with fewer than 500 genes per cell and cells with greater than 10% mitochondrial content. Clustering and batch effect were performed as has been previously described in these established methods^15^. Batch effects (as pertaining to day of capture) were removed by normalizing all cells in a batch to the most counts, and then multiplying by the median counts in that batch. These normalized counts were merged together across samples and log_2_ normalization was performed. Using default parameters of Seurat v2, variable genes were identified. Batch was regressed out in the space of variable genes during scaling, again using default Seurat v2 parameters. For three samples, there was only one individual per timepoint, which may have resulted in confounding age and batch, but because of the strong batch effects that result from 10X data, we moved ahead with the batch correction. In the space of the scaled, batch corrected variable genes, we performed PCA analysis. Significant PCs were carried forward, as identified using the calculation detailed in a prior publication. Using the RANN package, the top 10 nearest neighbors were identified for each cell in the space of significant principal components. We used a custom script to calculate the Jaccard distance of these neighbors, and used igraph to perform Louvain clustering for each analysis. Clustering with batch effect correction was performed for each dissected area, and also without batch correction by each individual, both across all brain regions and within cortex only.

Cluster markers were interpreted and assigned cluster identity by using known literature cell type annotations, or by associating progenitor and neuronal genes to other identifiable details. Progenitors were distinguished as radial glia if they expressed neurogenic genes, else they were determined to be neuroepithelial (STable 6).

Subclustering within the neurons was performed by selecting and clustering the cells from neuronal clusters and repeating the clustering procedure with batch effect correction. Progenitor cells were subclustered by removing neuronal, IPC, and other (microglia, endothelial, and pericyte) clusters from the data. After subclustering, neuronal populations were identified and removed iteratively until 54 subclusters were identified and did not include neuronal populations. These 54 subclusters were recombined by correlating marker genes to one another, and then performing a dendrogram cut to generate the 9 clusters presented here.

Correlational analysis between mouse and human data was performed as we previously described^7^. Briefly, cluster markers were generated for each dataset individually, and gene scores integrating the specificity and fold enrichment for each marker was calculated. A matrix for every marker gene and gene score was generated across all clusters and used for correlations. This was performed in the space of all mouse forebrain clusters as compared to primary human progenitor clusters (Fig 5), and in the space of all organoid clusters compared to the full set of primary human clusters in our dataset (SFig 23).

### RNA-velocity

Velocity estimates were calculated using the veloctyo.py v0.17 and scVelo (RRID:SCR_018168) algorithms. Reads that passed quality control after clustering were used as input for the velocyto command line implementation. The human expressed repeat annotation file was retrieved from the UCSC genome browser (RRID:SCR_005780). The genome annotation file used was provided via CellRanger. The output loom file was used as input to estimate velocity through scVelo. For each individual analysis, cells were filtered based on the following parameters: minimum total counts ≥ 200, minimum spliced counts ≥ 20 and minimum unspliced counts ≥ 10. For the combined cortical analysis, the processed loom files for each individual analysis were combined to generate a new UMI count matrix of 15,473 genes across 53,096 cells, for which the velocity embedding was estimated using the stochastic model. For the combined progenitor analysis, cells that were identified as progenitors were used to create the loom file. The loom files for each of the individuals were combined for a total count matrix of 14,207 genes across 30,562 cells for the velocity embedding using the same criteria. Each embedding was visualized using Uniform Manifold and Approximation and Projection of dimension reduction.

### Immunostaining

Primary human and mouse samples and organoids were collected, fixed in 4% paraformaldehyde, washed with 1x PBS and immersed in 30% sucrose in 1x PBS until saturated. Samples were embedded in cryomolds using 50% O.C.T. (Tissue-Tek Cat# 4583) and 50% of 30% sucrose in 1x PBS and frozen at -80°C. All samples were sectioned at 16μm onto SuperFrost Plus microscope slides. Citrate antigen retrieval (Vector Labs, Cat# H-3300) was performed for 20 min at ∼95-100°C. Slides were then washed with 1x PBS and blocked using 5% donkey serum, 2% gelatin, 0.1% Triton in 1x PBS for 90 min at room temperature (RT). Primary antibody incubation occurred in blocking buffer overnight at 4°C, and washed 5x using 0.3% Triton + 10mM glycine. Secondary antibody incubation occurred in blocking buffer for 2-3h at room temperature. Primary antibodies used: TRKC (1:200, R and D Systems Cat# AF373, RRID:AB_355332); ALX1 (1:500, Santa Cruz Biotechnology Cat# sc-130416, RRID:AB_2226324); ID4 (1:200, Santa Cruz Biotechnology Cat# sc-365656, RRID:AB_10859382); N-CADHERIN (1:300, Abcam Cat# ab18203, RRID:AB_444317); DLK1 (1:200, Abcam Cat# ab119930, RRID:AB_10902607); DLK1 (1:100, Abcam Cat# ab21682, RRID:AB_731965); CROC-4 (1:100, Aviva Systems Cat# ARP34802_P050, RRID:AB_2827813); LUM (1:50, Thermo Fisher Scientific Cat# MA5-29402, RRID:AB_2785270); ZO-1 (1:100, Thermo Fisher Scientific Cat# 61-7300, RRID:AB_2533938); SOX2 (1:100, R and D Systems Cat# AF2018, RRID:AB_355110); SOX2 (1:250, Santa Cruz Biotechnology Cat# sc-365823, RRID:AB_10842165); CTIP2 (1:500, Abcam Cat# ab18465, RRID:AB_2064130); KI67 (1:200, Thermo Fisher Scientific Cat# 14-5698, RRID: AB_10854564); HOPX (1:250, Santa Cruz Biotechnology Cat# sc-398703, RRID:AB_2687966); HOPX (1:200, Proteintech Cat# 11419-1-AP, RRID:AB_10693525); TBR2 (1:250, Abcam Cat# ab23345, RRID:AB_778267); TBR2 (1:250, R and D Systems Cat# AF6166, RRID:AB_10569705); NESTIN (1:200, Millipore Cat# MAB5326, RRID:AB_2251134), DCX (1:500, Aves Labs Cat# DCX, RRID:AB_2313540); NEUN (1:250, Millipore Cat# ABN91, RRID:AB_11205760); PAX6 (1:200, BioLegend Cat# 901301, RRID:AB_2565003); FOXG1 (1:1000, Abcam Cat# ab18259, RRID:AB_732415); SATB2 (1:250, Abcam Cat# Ab51502, RRID:AB_882455); LHX5 (1:100, R and D Systems Cat# AF6290, RRID:AB_10973257); REELIN (1:100, MBL Cat# D223-3, RRID:AB_843523); Phospho-β-CATENIN (1:100, Cell Signaling Technology Cat# 9561, RRID:AB_331729); Phospho-S6 (1:100, Cell Signaling Technology Cat# 2211S, RRID:AB_331679); NICD/NOTCH1 (1:100, Millipore Cat# 07-1232, RRID:AB_1977387). All secondary antibodies were AlexaFluor used at a dilution 1:1000. Secondary antibodies: Donkey anti-Mouse 488 (Thermo Fisher Scientific Cat# A32766, RRID:AB_2762823); Donkey anti-Rabbit 488 (Thermo Fisher Scientific Cat# A32790, RRID:AB_2762833); Donkey anti-Chicken 488 (Jackson ImmunoResearch Labs Cat# 703-545-155, RRID:AB_2340375); Donkey anti-chicken 594 (Jackson ImmunoResearch Labs Cat# 703-585-155, RRID:AB_2340377); Donkey anti-Mouse 546 (Thermo Fisher Scientific Cat# A10036, RRID:AB_2534012); Donkey anti-Mouse 594 (Thermo Fisher Scientific Cat# A-21203, RRID:AB_141633); Donkey anti-Mouse 647 (Thermo Fisher Scientific Cat# A32787, RRID:AB_2762830); Donkey anti-Mouse 680 (Thermo Fisher Scientific Cat# A32788, RRID:AB_2762831); Donkey anti-Rabbit 546 (Thermo Fisher Scientific Cat# A10040, RRID:AB_2534016); Donkey anti-Rabbit 594 (Thermo Fisher Scientific Cat# A-21207, RRID:AB_141637); Donkey anti-Rabbit 647 (Thermo Fisher Scientific Cat# A32795, RRID:AB_2762835); Donkey anti-Rat 594 (Thermo Fisher Scientific Cat# A-21209, RRID:AB_2535795); Donkey anti-Rat 488 (Thermo Fisher Scientific Cat# A-21208, RRID:AB_2535794); Donkey anti-Goat 546 (Thermo Fisher Scientific Cat# A-11056, RRID:AB_2534103); Donkey anti-Goat 594 (Thermo Fisher Scientific Cat# A-11058, RRID:AB_2534105); Donkey anti-Goat 647 (Thermo Fisher Scientific Cat# A32849, RRID:AB_2762840); Donkey anti-Sheep 546 (Thermo Fisher Scientific Cat# A-21098, RRID:AB_2535752); Donkey anti-Sheep 594 (Thermo Fisher Scientific Cat# A-11016, RRID:AB_2534083); Donkey anti-Sheep 647 (Thermo Fisher Scientific Cat# A-21448, RRID:AB_2535865); Donkey anti-Guinea Pig 647 (Jackson ImmunoResearch Labs Cat# 706-605-148, RRID:AB_2340476).

### In situ hybridization

Primary fixed samples were treated using the protocol for RNAScope Multiplex Fluorescence Assay v2 (Advanced Cell Diagnostics Cat# 323100) for C1orf61 amplification, targeting nucleotides 60-897 of NM_006365.3 (Advanced Cell Diagnostics Probe Design # NPR-0003991); MEF2C amplification, targeting nucleotides 1058 – 2575 of NM_002397.4 (Advanced Cell Diagnostics Cat# 452881); and the protocol for BaseScope v2 Assay (Advanced Cell Diagnostics Cat# 323900) for chromogenic LHX5-AS1 amplification, targeting nucleotides 69-293 of NR_126425.1 (Advanced Cell Diagnostics Probe Design # NPR-0003991).

### Imaging and Image Processing

Images were collected on the Leica SP8 (RRID:SCR_018169) inverted confocal microscope using a 40X oil-immersion objective. Because of the scarcity of first trimester primary tissue, only one sample per panel was imaged. For each imaging panel, the parameters (including the gain, offset, pinhole and laser power) for image acquisition was left constant for all samples. Images were later processed using FIJI Image J (RRID:SCR_003070).

### Statistical Tests

No statistical methods were used to pre-determine sample sizes. No randomization was used in this study. Distributions of the data were nor tested. Data collection and analysis were not performed blind to the conditions of the experiments. The Wilcoxon rank sum test was used to calculate clustermarkers within Seurat for a variety of analyses. A one-sided t-tested was used in SFig 1E and SFig 15C. A loess regression was used to estimate smoothed gene expression in SFig 11 and Figure 4.

### Code Availability

No custom code was used in this study. Open source algorithms were used as detailed in Single-cell analysis methods. However, any details on how these algorithms were used are available from the corresponding author upon request.

## Data Availability

The data that support the findings of this study are available from the corresponding author upon request. Raw single-cell sequencing data is available from the NeMO Repository at https://assets.nemoarchive.org/dat-0rsydy7. Processed single-cell sequencing data and full tilescan images are available for exploration and for download at our cell browser: https://cells-test.gi.ucsc.edu/?ds=early-brain.

## Supplementary Table Legends

**Supplementary Table 1. Whole Brain Metadata** Each cell that was used in this study is presented in this table with information about number of genes, number of UMIs, sample identity, sample age, cluster identity based upon whole brain clustering (though this analysis had some batch effect and was not used in this study), area, percent ribosomal content, area as annotated during actual dissection, and the cluster the cell belongs to based upon individual clusterings.

**Supplementary Table 2. Whole Brain Clustering By Individual Areas** Each of the 10 dissected areas was clustered across all samples with the same dissection; batch correction was used to mix the samples with one another. The resulting cluster markers as calculated by the Wilcoxon rank test are presented here for each area, including the p-value, the average log_2_(fold change) of the gene in the cluster compared to all other clusters, the percent of cells expressing the gene in the cluster (pct.1), the percent of cells expressing the gene in all other clusters (pct.2), and the adjusted p-value.

**Supplementary Table 3. Whole Brain Clustering By Individual** Each of the 9 individuals were clustered across all dissections. The resulting cluster markers as calculated by the Wilcoxon rank test are presented here for each area, including the p-value, the average log_2_(fold change) of the gene in the cluster compared to all other clusters, the percent of cells expressing the gene in the cluster (pct.1), the percent of cells expressing the gene in all other clusters (pct.2), and the adjusted p-value.

**Supplementary Table 4. Differential Gene Expression Across Areas Within a Single Individual** In each individual dataset, differential gene expression (Wilcoxon rank test) was performed across all dissected brain regions. The results from the oldest samples (CS22 and CS22_2) were used to identify the markers for each area and subsequently assessed across developmental time (SFig 1 – 2). The resulting cluster markers as calculated by the Wilcoxon rank test are presented here for each area, including the p-value, the average log_2_(fold change) of the gene in the cluster compared to all other clusters, the percent of cells expressing the gene in the cluster (pct.1), the percent of cells expressing the gene in all other clusters (pct.2), and the adjusted p-value.

**Supplementary Table 5. Cortex Annotations** Additional metadata similar to STable 1 is present for the cortex samples only. This also includes the progenitor cluster annotation.

**Supplementary Table 6. Cortex Clustermarkers** Clustering was performed across all cortical cells, resulting in 61 clusters that were subsequently annotated by cell type. The resulting cluster markers as calculated by the Wilcoxon rank test are presented here for each area, including the p-value, the average log_2_(fold change) of the gene in the cluster compared to all other clusters, the percent of cells expressing the gene in the cluster (pct.1), the percent of cells expressing the gene in all other clusters (pct.2), and the adjusted p-value.

**Supplementary Table 7. Cortical Neuron Subclustering** Neurons were subclustered and cluster markers were calculated. Neurons from early samples were similarly subclustered and presumed to result from direct neurogenesis. Within the neurons, differential expression was performed between older and younger neuronal populations across all subtypes. The resulting cluster markers for these three analysis are presented as calculated by the Wilcoxon rank test are presented here for each area, including the p-value, the average log_2_(fold change) of the gene in the cluster compared to all other clusters, the percent of cells expressing the gene in the cluster (pct.1), the percent of cells expressing the gene in all other clusters (pct.2), and the adjusted p-value.

**Supplementary Table 8. Progenitor Subclustered Cluster Markers** Progenitors were subclustered and cluster markers were calculated. The resulting cluster markers for these three analysis are presented as calculated by the Wilcoxon rank test are presented here for each area, including the p-value, the average log_2_(fold change) of the gene in the cluster compared to all other clusters, the percent of cells expressing the gene in the cluster (pct.1), the percent of cells expressing the gene in all other clusters (pct.2), and the adjusted p-value.

**Supplementary Table 9. Ultra Early Cortex Cluster Markers** The earliest cortical samples (CS12, CS13) in the dataset were subclustered to look for unique populations within these least characterized timepoints. The resulting cluster markers for this analysis are presented as calculated by the Wilcoxon rank test are presented here for each area, including the p-value, the average log_2_(fold change) of the gene in the cluster compared to all other clusters, the percent of cells expressing the gene in the cluster (pct.1), the percent of cells expressing the gene in all other clusters (pct.2), and the adjusted p-value.

**Supplementary Table 10. Velocity Genes and Disease Relevance** Velocity genes were calculated using default parameters, and grouped by cluster. Genes were ranked based on velocity score, and compared across all cortical cells, progenitor cells and the cells for each individual sample. Genes that appear in ≥ 3 individual, cortical or progenitor samples were explored for their functional and clinical significance. Genes associated with neurodevelopmental disorders are listed in this table with their corresponding PMID references.

**Supplementary Table 11. Early Versus Late Differential Gene Expression** Differential gene expression was calculated between the early samples in the dataset (CS12-15) and the later samples in the dataset (CS19-22). The resulting cluster markers for this analysis are presented as calculated by the Wilcoxon rank test are presented here for each area, including the p-value, the average log_2_(fold change) of the gene in the cluster compared to all other clusters, the percent of cells expressing the gene in the cluster (pct.1), the percent of cells expressing the gene in all other clusters (pct.2), and the adjusted p-value. These lists were then compared to the CellPhoneDB curated interactions list, and two potential ligand/receptor interactions were identified (NTRK3 and DLK1) and bolded within the table.

**Supplementary Table 12. Progenitor WGCNA** Weighted Gene Coexpression Analysis (WGCNA) was performed using the same parameters are previous single-cell studies (top 5000 loaded genes from significant PCs), dendrogram cut at 4. This yielded 98 networks, and genes are in column A and the network names are in column B.

## Notes

https://assets.nemoarchive.org/dat-0rsydy7

https://cells-test.gi.ucsc.edu/?ds=early-brain

### Competing Interest Statement

The authors have declared no competing interest.

