## Supplemental Figures for "Single-Cell Atlas of Early Human Brain Development Highlights Heterogeneity of Human Neuroepithelial Cells and Early Radial Glia"

### Supplementary Figures and Legends

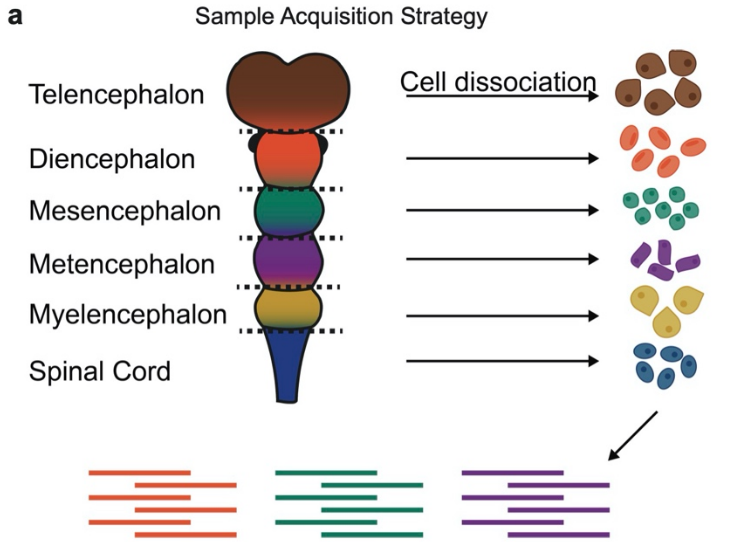


**b** Dataset Summary By Area

| Area | Number of Cells | Number of Individuals |
|---|---|---|
| Cortex | 58145 | 10 |
| Ganglionic Eminences | 24835 | 6 |
| Retina | 8978 | 4 |
| Olfactory Placode | 2040 | 1 |
| Thalamus | 24091 | 4 |
| Hypothalamus | 13275 | 3 |
| Midbrain | 31402 | 7 |
| Hindbrain | 60000 | 8 |
| Cerebellum | 14769 | 2 |
| Spinal Cord | 11754 | 1 |

**c** Dataset Summary By Individual

| Individual | Number of Cells | Number of Areas |
|---|---|---|
| CS12 | 28412 | 6 |
| CS13 | 31462 | 6 |
| CS14 | 4633 | 3 |
| CS14_3 | 31250 | 4 |
| CS15 | 166 | 3 |
| CS15_2 | 30566 | 4 |
| CS19 | 22215 | 4 |
| CS20 | 53214 | 7 |
| CS22 | 23414 | 8 |
| CS22_2 | 23945 | 8 |

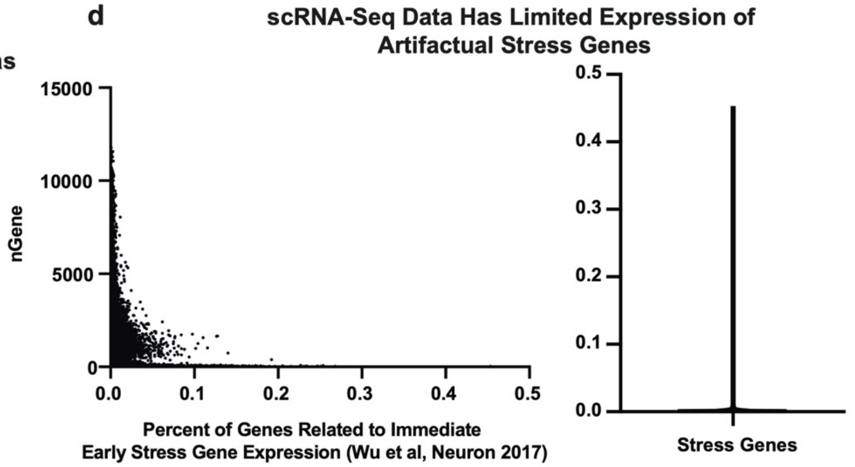


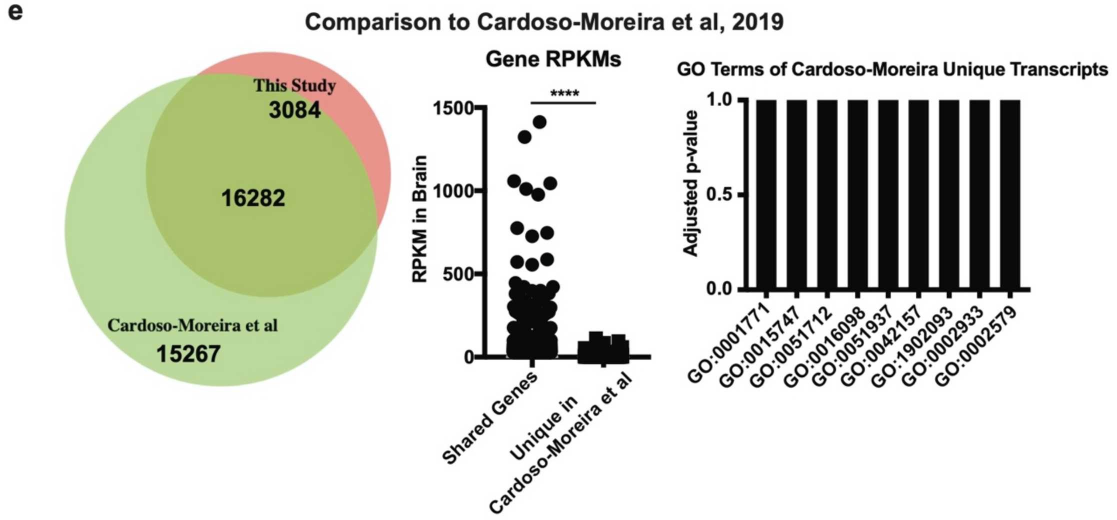


**Supplementary Figure 1. Sample Acquisition and Quality Control a)** Samples were obtained from the HDBR and dissected into anatomical regions. These regions were then dissociated into a single-cell suspension and captured with 10X Chromium v2. **b)** Summary of dataset collected by region, number of cells passing quality control per region, and the number of individuals that were collected for each region. **c)** Summary of dataset collected by individual, number of cells passing quality control per region, and the number of areas that were collected for each individual. **d)** Using the gene lists compiled in Wu et al, Neuron, 2017 we examined our single-cell RNA-sequencing data for early stress gene expression which is a common artefactual activation state for dissociated cells. We observed that cells expressing these gene signatures was minimal, and those that had high expression of this stress gene set also had a low number of genes/cell and were eliminated from further analysis for this reason. **e)** We compared the expression of genes in our data to the bulk RNA-sequencing at equivalent timepoints from Cardoso-Moreira et al, 2019. We find substantial overlap, and also find that the RPKM of shared genes (n = 16,282 genes) is significantly higher than that of those in the bulk dataset alone (n = 15,267 genes) (One sided t-test, p-value = $1.467e^{-194}$).

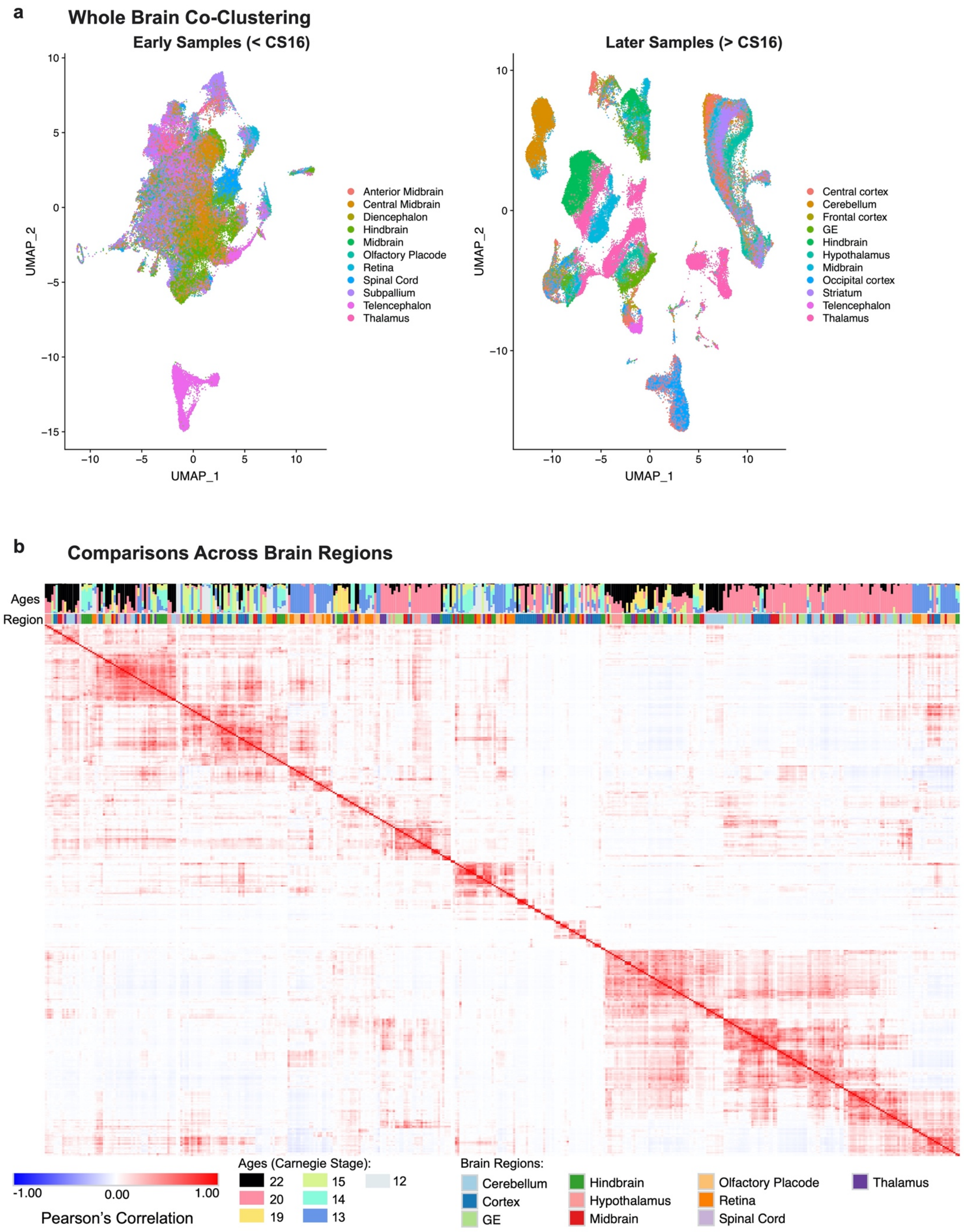
a
Whole Brain Co-Clustering
Early Samples (< CS16)
Later Samples (> CS16)
UMAP_1
UMAP_2
Anterior Midbrain
Central Midbrain
Diencephalon
Hindbrain
Midbrain
Olfactory Placode
Retina
Spinal Cord
Subpallium
Telencephalon
Thalamus
Central cortex
Cerebellum
Frontal cortex
GE
Hindbrain
Hypothalamus
Midbrain
Occipital cortex
Striatum
Telencephalon
Thalamus
b
Comparisons Across Brain Regions
Ages
Region
-1.00
0.00
1.00
Pearson's Correlation
Ages (Carnegie Stage):
22
20
19
15
14
13
12
Brain Regions:
Cerebellum
Cortex
GE
Hindbrain
Hypothalamus
Midbrain
Olfactory Placode
Retina
Spinal Cord
Thalamus

**Supplementary Figure 2. Whole Brain Clustering and Cell Types Across Brain Regions a)** Clustering was performed across all brain structures in early and late samples. UMAP plots are shown colored by the sample area of origin. At early stages, samples do not segregate by their area of origin, likely because their stemness programs are more similar than areal specification signals are different. At later stages, samples begin to segregate by brain region of origin. **b)** Clusters were identified in each individual clustering across the 10 regions shown in the legend. These clusters were correlated using a Pearson's Correlation to one another as described in Figure 4 and Methods. Hierarchical clustering of these correlations identified two major groups (left: progenitors; right: neurons) that showed that the cell type programs were identifiable across brain regions (shown above the heatmap) and ages (graph above heatmap shows representative fraction of cells in given cluster from each of the ages, legend at bottom).

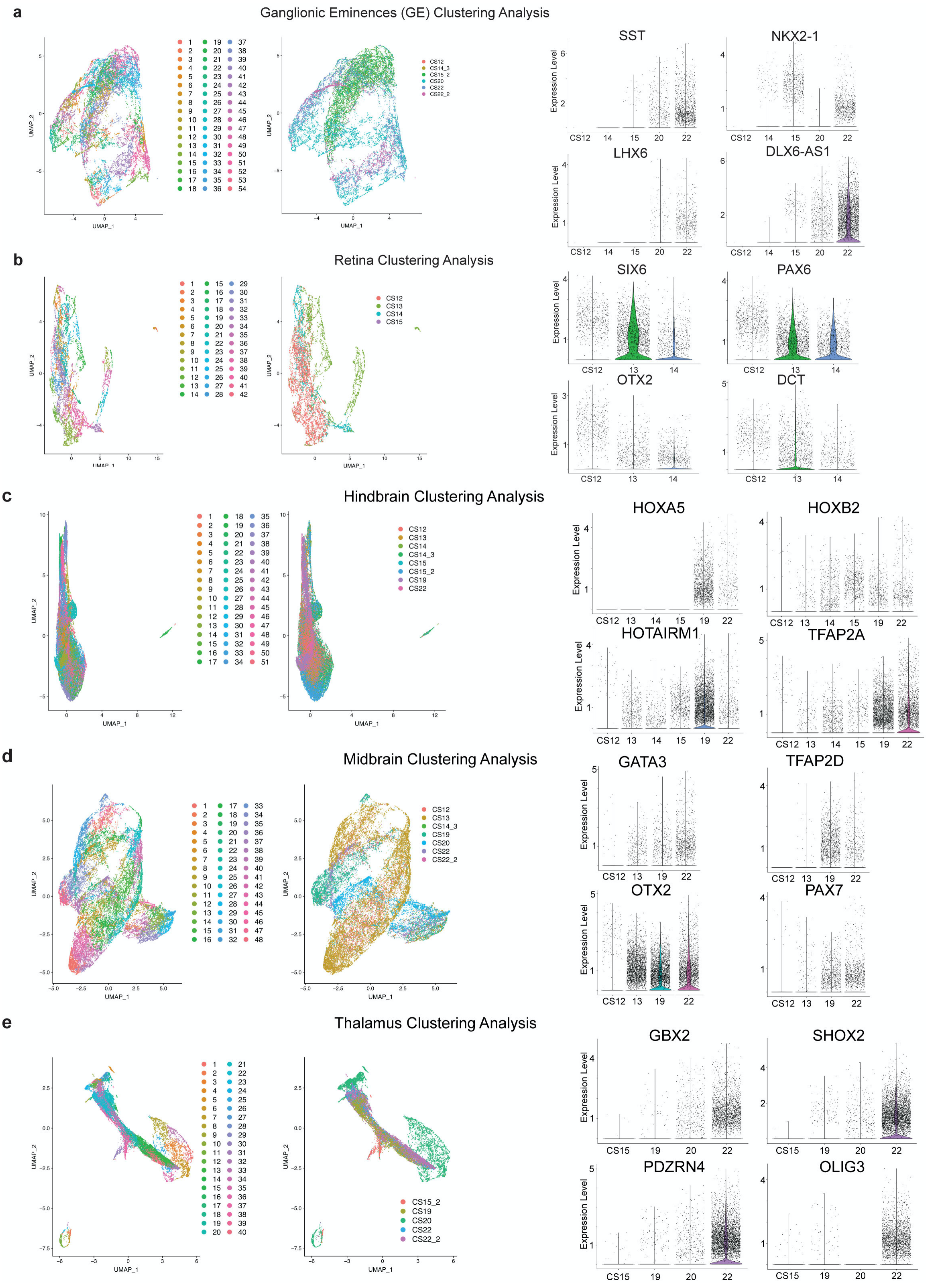

a
Ganglionic Eminences (GE) Clustering Analysis
SST
NKX2-1
LHX6
DLX6-AS1
b
Retina Clustering Analysis
SIX6
PAX6
OTX2
DCT
c
Hindbrain Clustering Analysis
HOXA5
HOXB2
HOTAIRM1
TFAP2A
d
Midbrain Clustering Analysis
GATA3
TFAP2D
OTX2
PAX7
e
Thalamus Clustering Analysis
GBX2
SHOX2
PDZRN4
OLIG3
Expression Level
UMAP_1
UMAP_2

**Supplementary Figure 3. Clustering by Brain Regions a)** Region specific clustering of ganglionic eminence samples. UMAP on the left has the clusters, and the UMAP on the right has the individuals from which these samples were collected. On the far right, violin plots show 4 highly region-specific genes as identified from area-specific differential expression in the oldest samples, with their emergence across ages shown in the violin plots. **b)** Region specific clustering of retina samples with same layout as is found in (a). **c)** Region specific clustering of hindbrain samples with same layout as is found in (a). **d)** Region specific clustering of midbrain samples with same layout as is found in (a).

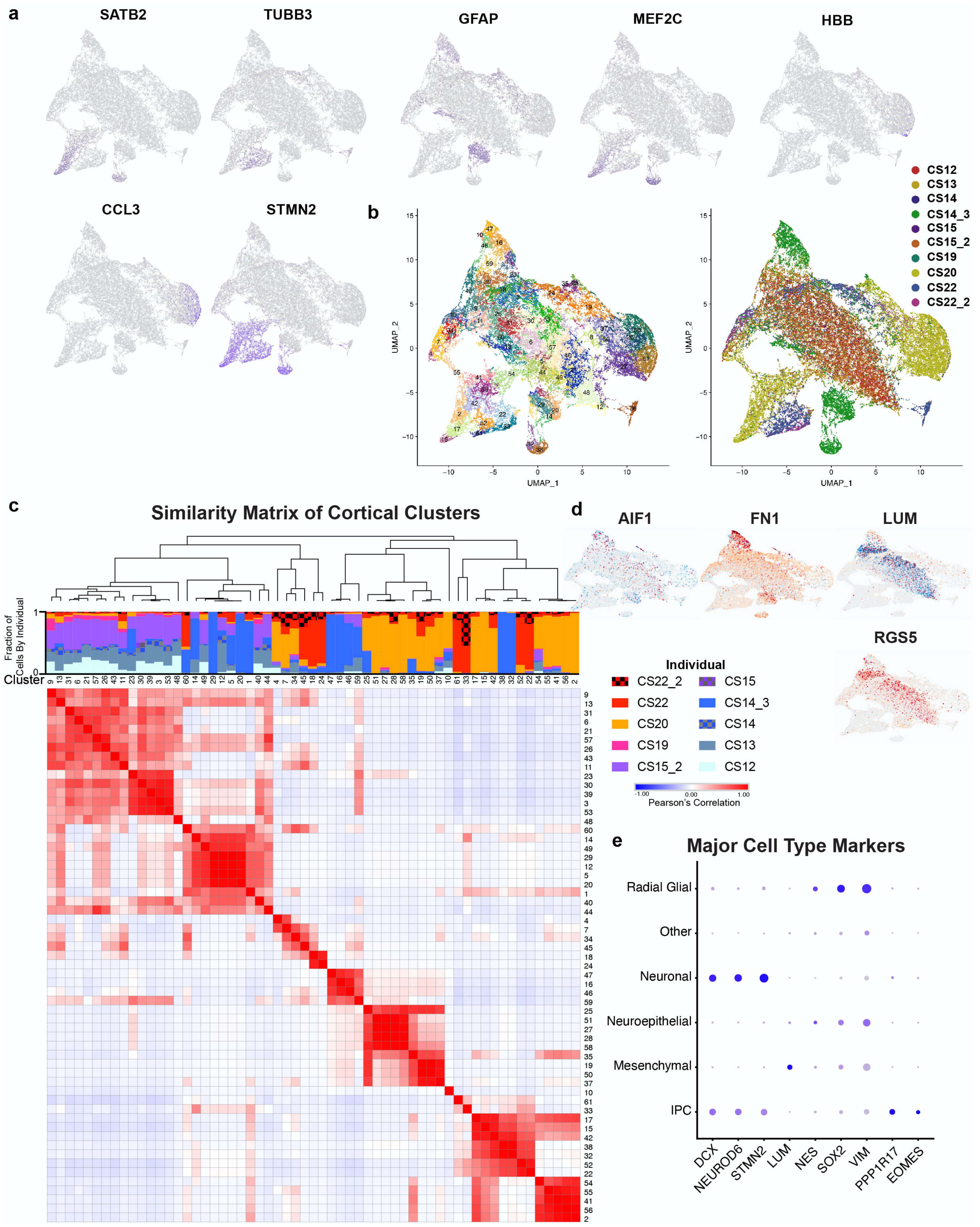


**Supplementary Figure 4. Additional Characterization of Cortical Cell Types a)** Feature plots of additional marker genes including neuronal markers (*SATB2, TUBB3, MEF2C, STMN2*), maturing progenitors (*GFAP*), red blood cells (*HBB*), and microglia (*CCL3*). **b)** UMAP on the left shows the 61 clusters that have been identified in the early developing cortex and designated by cell type for Figure 1. The cluster markers and annotations are presented in STable 6. The UMAP on the right shows the clustering by individual, with each individual marked by age and additional markings for multiple individuals of the same age. **c)** Hierarchical clustering of similarity scores across all cortical clusters, with intensity of similarity indicated from -1 to 1 by a blue to red scale. For each cluster, the composition by individual is shown as a stacked barplot at the top of the heatmap, with the majority of clusters containing contributions from multiple individuals. **d)** Velocity plots showing dynamics of relative gene expression for *AIF1*, *FN1*, *LUM*, and *RGS5*. **e)** Dot plot of major cell type markers across annotated cell types.

a

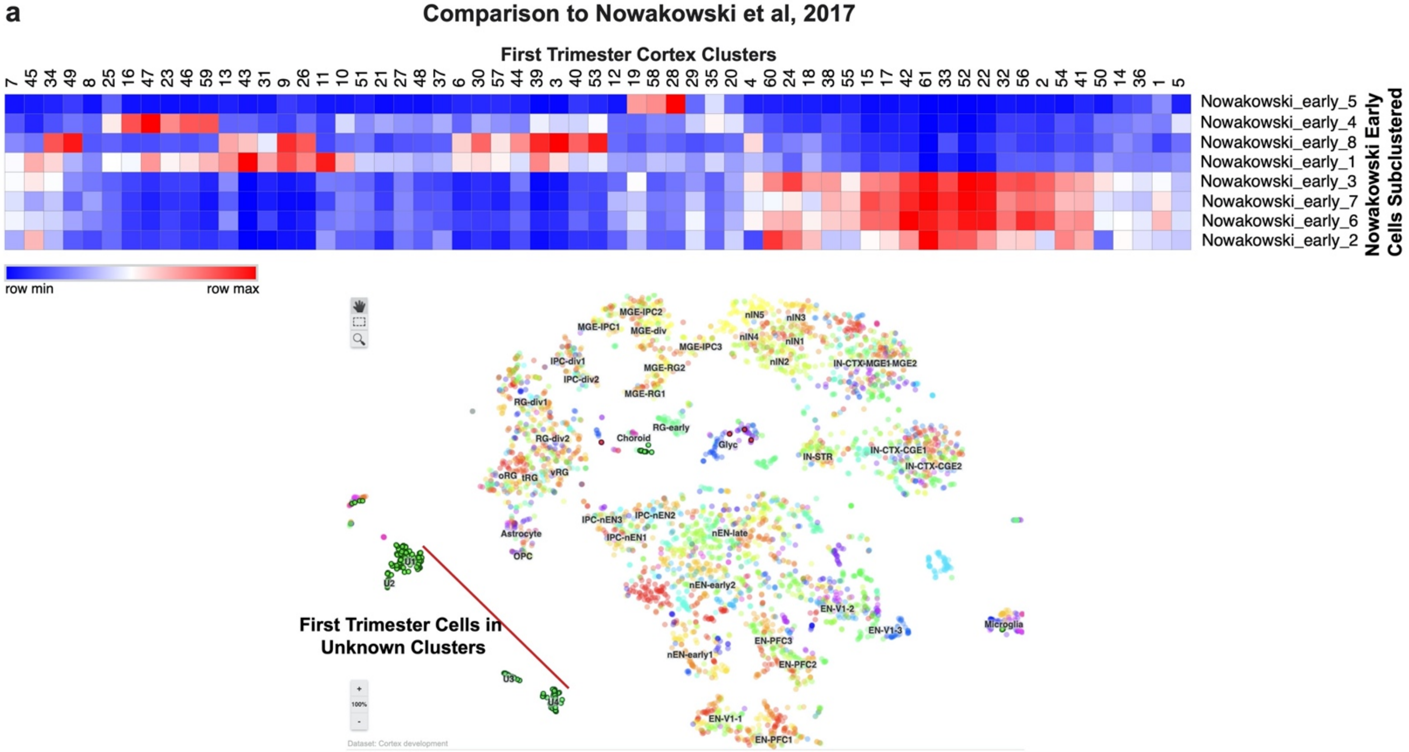


b

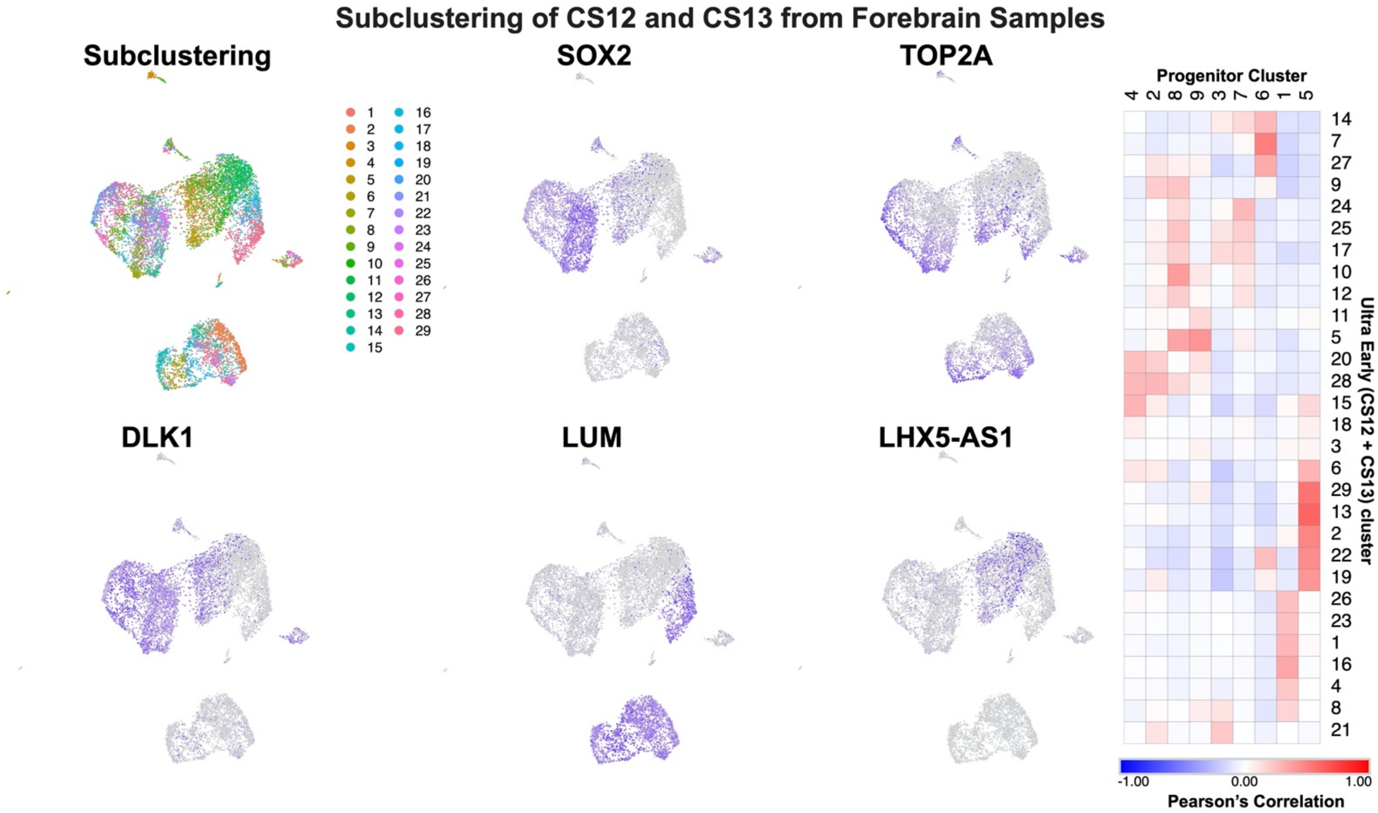


**Supplementary Figure 5. Comparison of Early Cell Populations a)** We compared our clusters to the first trimester cells captured in Nowakowski et al, Science, 2017. First, we analyzed the early cells from the Nowakowski dataset and identified 8 sub-clusters. Correlation of these clusters to our 63 clusters identified strong correspondence (> 0.4) for all 8 clusters, highlighting that using these correlations we could interpret the cell types from the Nowakowski data. As shown in the screenshot from the annotated cell browser below, in the previous analysis of these cell types, these first trimester cells were annotated to have an "unknown" identity. **b)** To better understand the heterogeneity of the earliest samples in the data, we subclustered the CS12 and CS13 telencephalon cells independently of the remaining data. We identified 29 subclusters that primarily represented 4 groups marked by SOX2-positive/DLK1-positive progenitor cells, TOP2A-positive dividing cells, LUM-positive mesenchymal cells, and LHX5-AS1 positive cells.

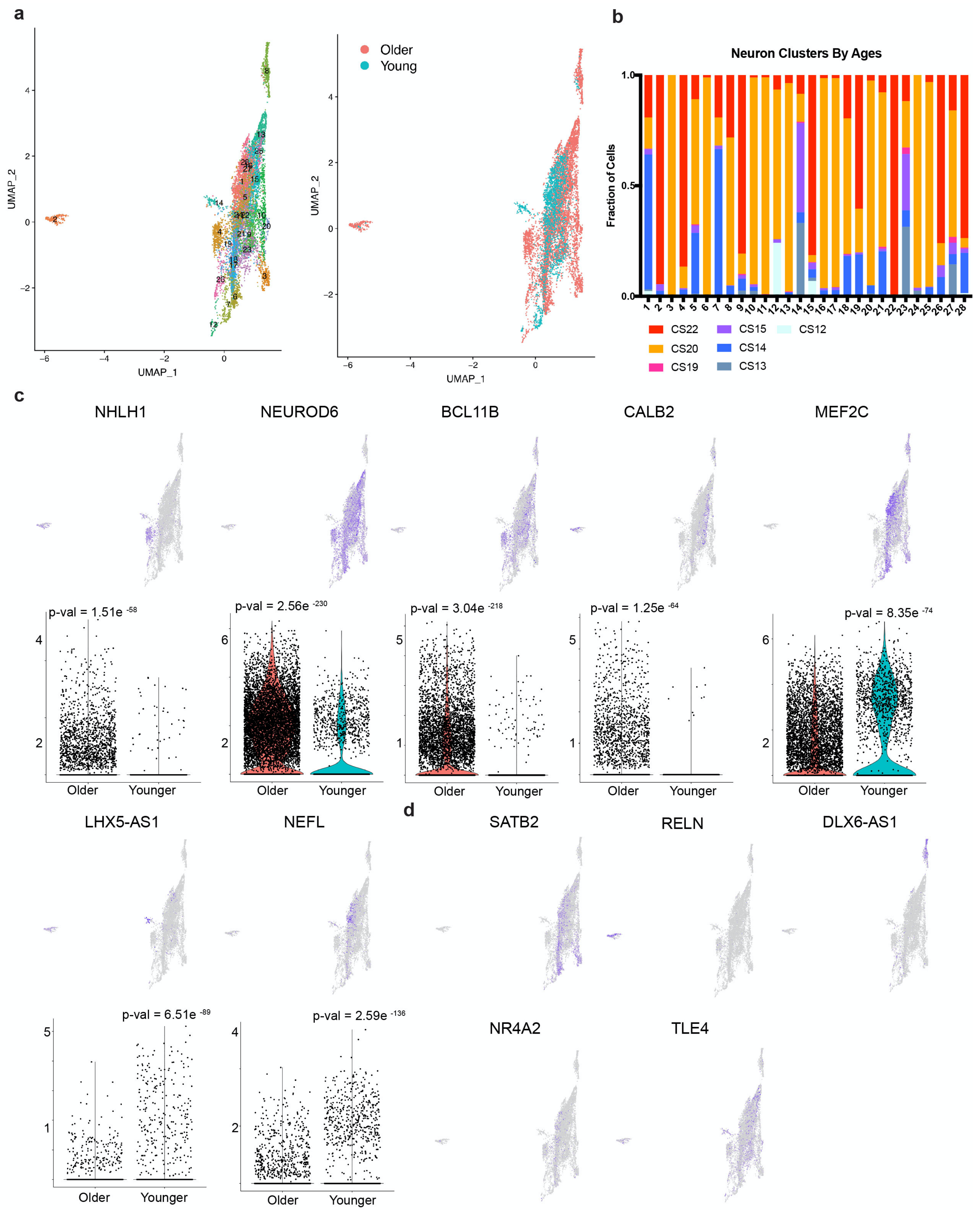


**Supplementary Figure 6. Sub-clustering of Neuronal Populations Identifies Distinct Neurons from Direct and Indirect Neurogenesis a)** UMAP on the left depicts the clusters identified from the subclustering of all neuronal populations identified in the cortical samples. UMAP on the right depicts the segregation of these clusters by younger samples (CS12-15, before IPCs are identifiable, presumably from direct neurogenesis) and older samples (CS19-22, after IPCs are identifiable and presumably from indirect neurogenesis). **b)** Composition of neuronal subclusters by sample ages. Some clusters are well mixed between older and younger samples, but most are specific to one group or another. **c)** Feature plots are shown for cluster markers for clusters in (a) (see also STable X) that also have significant differences between older and younger samples, with violin plots below showing the relative distribution in older and younger samples with p-values written above (Wilcoxon rank sum test, test is not sided but is corrected for multiple hypothesis testing). **d)** Additional marker gene feature plots are shown that do not have significant differences between older and younger samples (Wilcoxon rank sum test, test is not sided but is corrected for multiple hypothesis testing). *SATB2* is typically a marker of upper layer neurons generated at later stages of development, so it is unexpected to see expression at the early time points. *RELN* marks Cajal-Retzius (CR) cells that are thought to be some of the earliest born, and we see a few cells from the early samples and many CR cells in the older samples. *DLX6-AS1* marks inhibitory interneurons that migrate in from the ventral telencephalon and are primarily comprised of cells from older samples, as would be expected. *TLE4* and *NR4A2* are markers of subplate and appear to mark distinct clusters in both younger and older samples, hinting towards additional heterogeneity of subplate neurons.

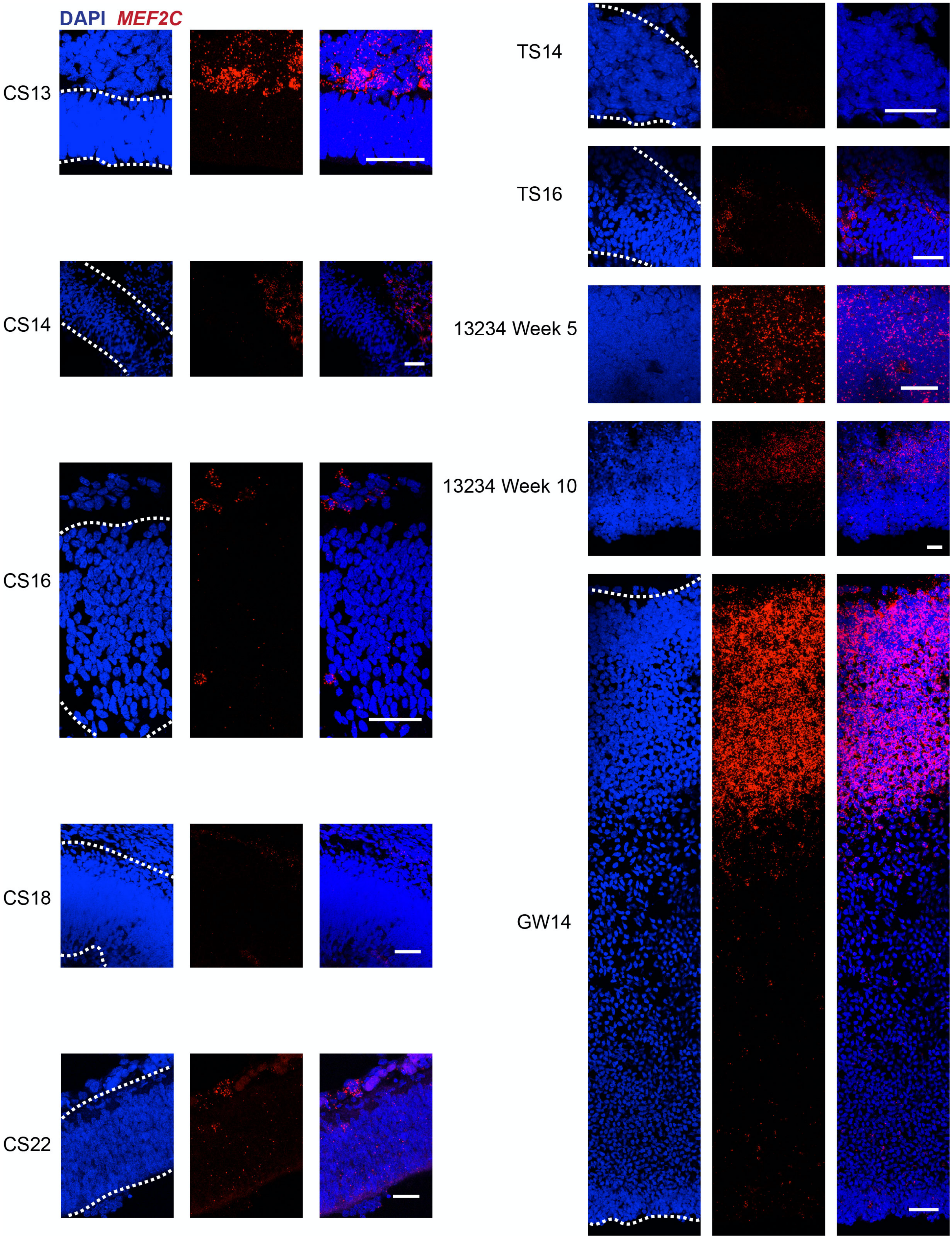
DAPI MEF2C
CS13
CS14
CS16
CS18
CS22
TS14
TS16
13234 Week 5
13234 Week 10
GW14

**Supplementary Figure 7. MEF2C RNA Expression in the Developing Human Brain** *MEF2C* marks a subset of the neuronal populations but surprisingly also Progenitor Cluster 1. *MEF2C* (red) RNA expression via fluorescent *in situ* hybridization marks a large extracortical population of cells near the developing cortical plate at CS13, 14 and 16 that begins to dissipate by CS22. By GW14 its expression is much higher and also only marks neurons. There is no *MEF2C* expression in the mouse samples until TS16. However, there is diffuse expression for *MEF2C* in the cerebral organoid sample (13234) at Week 5 that becomes less abundant and more localized by Week 10. Nuclei are marked by DAPI in blue. Each sample was processed and stained one time. All scale bars are 50 μM.

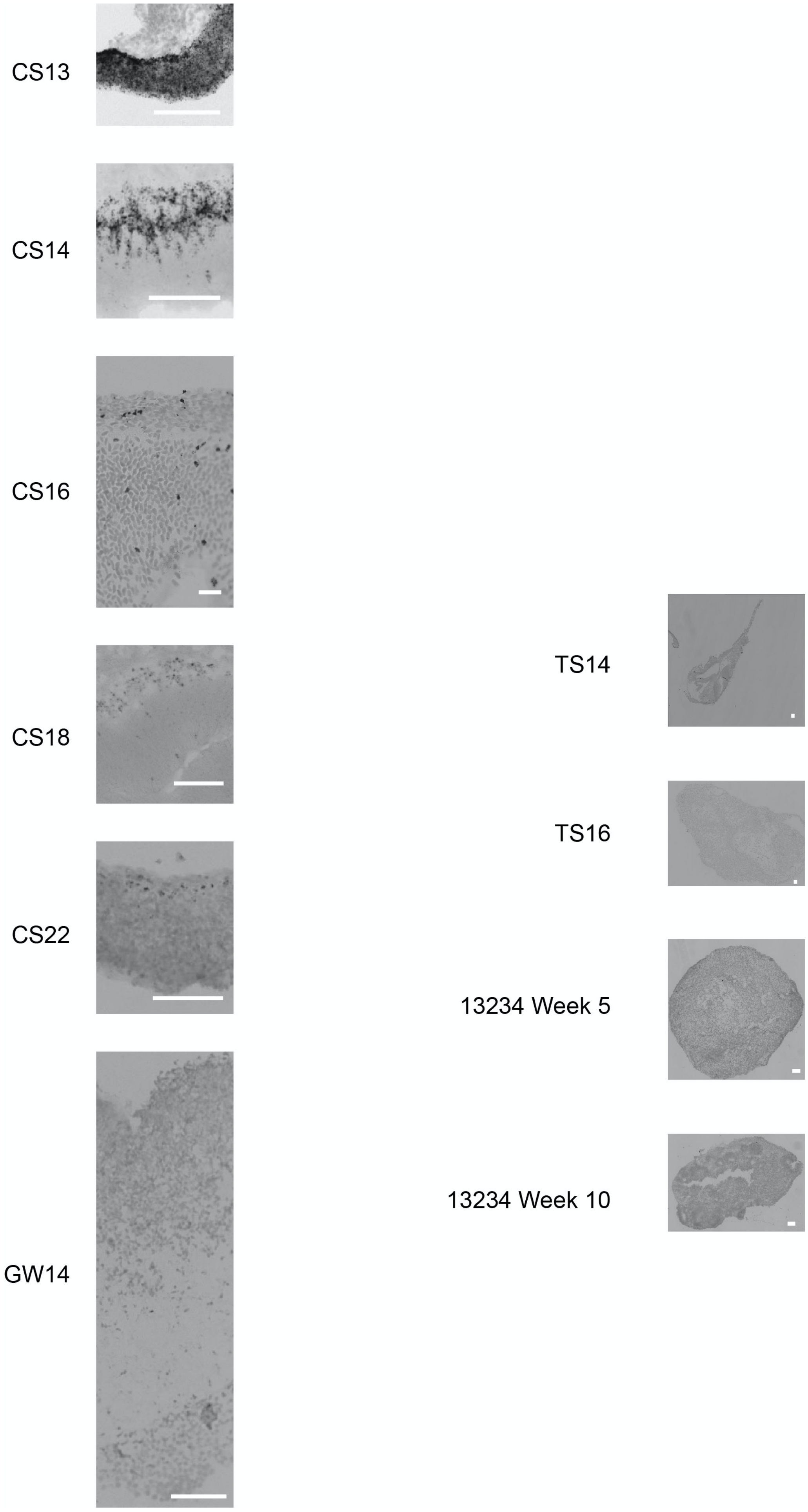
CS13
CS14
CS16
CS18
CS22
GW14
TS14
TS16
13234 Week 5
13234 Week 10

**Supplementary Figure 8. *LHX5-AS1* Expression in the Developing Human Brain** *LHX5-AS1* marks Progenitor Cluster 8. Chromogenic *in situ* hybridization demonstrates a pattern of RNA expression that mimics that of LHX5 (SFig 21). *LHX5-AS1* is diffusely present at CS13, but by CS14 its expression is limited to the developing cortical plate. By GW14, there is no longer any *LHX5-AS1* expression. There is no *LHX5-AS1* expression in the mouse samples at either TS14 or TS16, and there was very limited expression in the organoid (13234) at either stage (Week 5 or Week 10). Each sample was processed and stained one time. All scale bars are 50 μM.

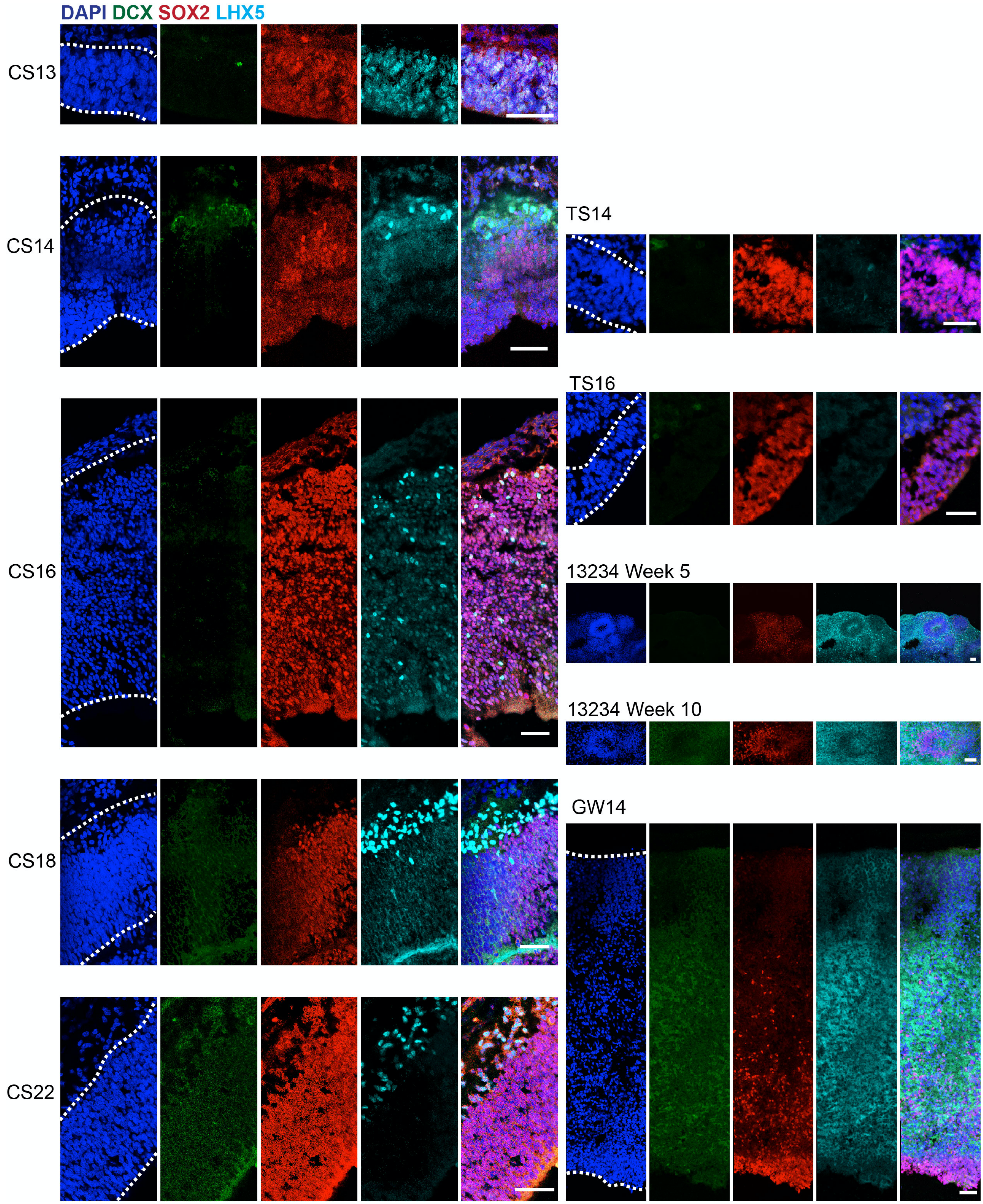


**Supplementary Figure 9. LHX5 Expression in the Developing Human Brain** LHX5 marks Progenitor Cluster 8. Immunostaining demonstrates nearly ubiquitous staining of LHX5 in the CS13 cortical sample. However, by CS14 its staining is limited to the developing cortical plate (DCX, green). By GW14, there is only background staining (no presumptive nuclei present). There is very minimal staining of LHX5 in the mouse sample at TS14 that abolishes by TS16. There is abundant staining in the organoid samples (13234) at both stages (Week 5 and Week 10). Progenitors are marked by SOX2 in red. Nuclei are marked by DAPI in blue. Each sample was immunostained one time. All scale bars are 50 μM.

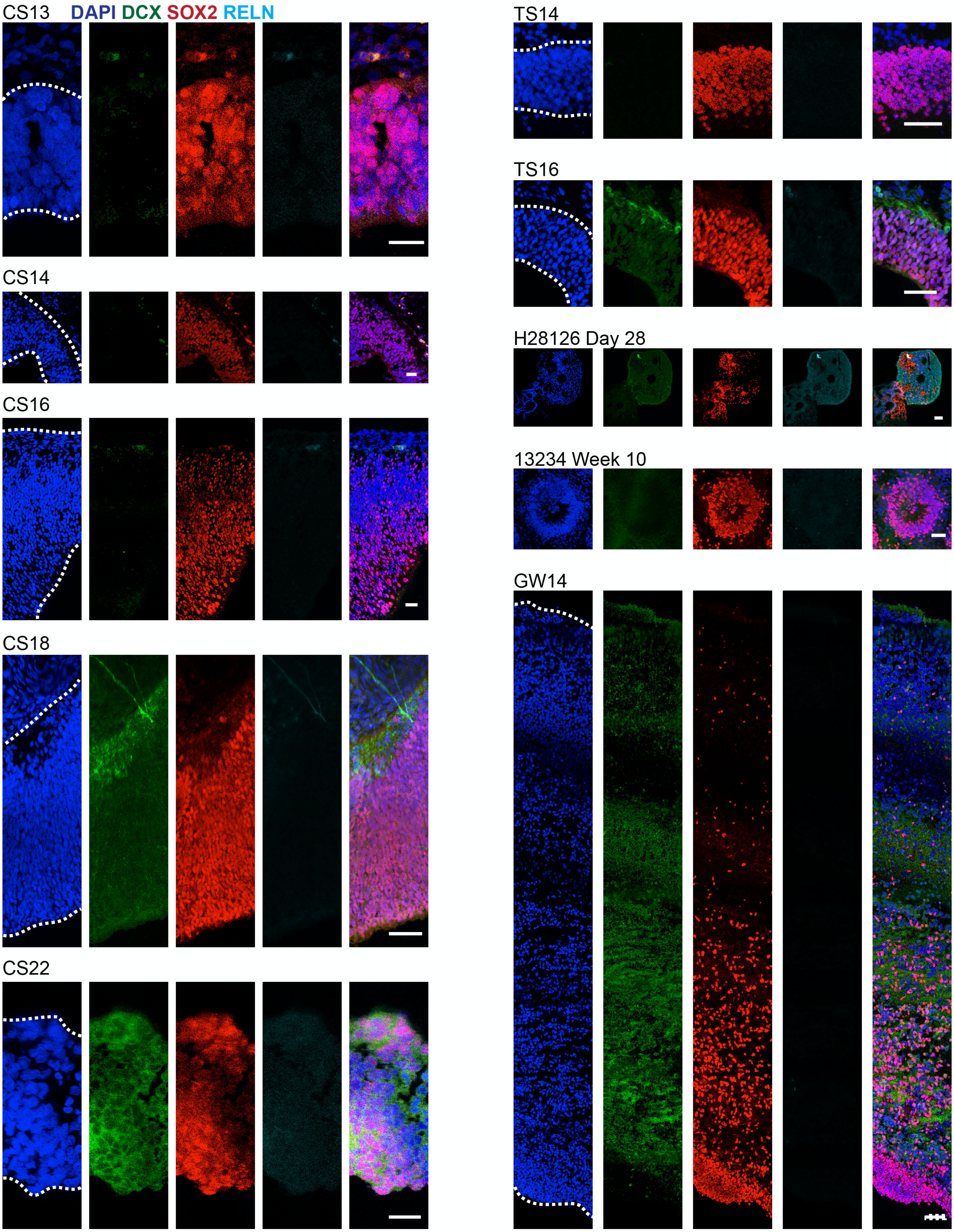

CS13
DAPI DCX SOX2 RELN
CS14
CS16
CS18
CS22
TS14
TS16
H28126 Day 28
13234 Week 10
GW14

**Supplementary Figure 10. RELN Expression in the Developing Human Brain** RELN is a marker for Cajal-Retzius cells and is identified in the neuronal population of this dataset. RELN (cyan) only marks this neuronal population (DCX, green) in the earliest samples (CS13 and CS14). By CS22, RELN staining is no longer present. Immunostaining for RELN shows very minimal staining in the TS16 mouse sample. Cerebral organoid samples (H28126 and 13234) also show minimal immunostaining for RELN. Progenitors are marked by SOX2 in red. Nuclei are marked by DAPI in blue. Each sample was immunostained one time. All scale bars are 50 μM.

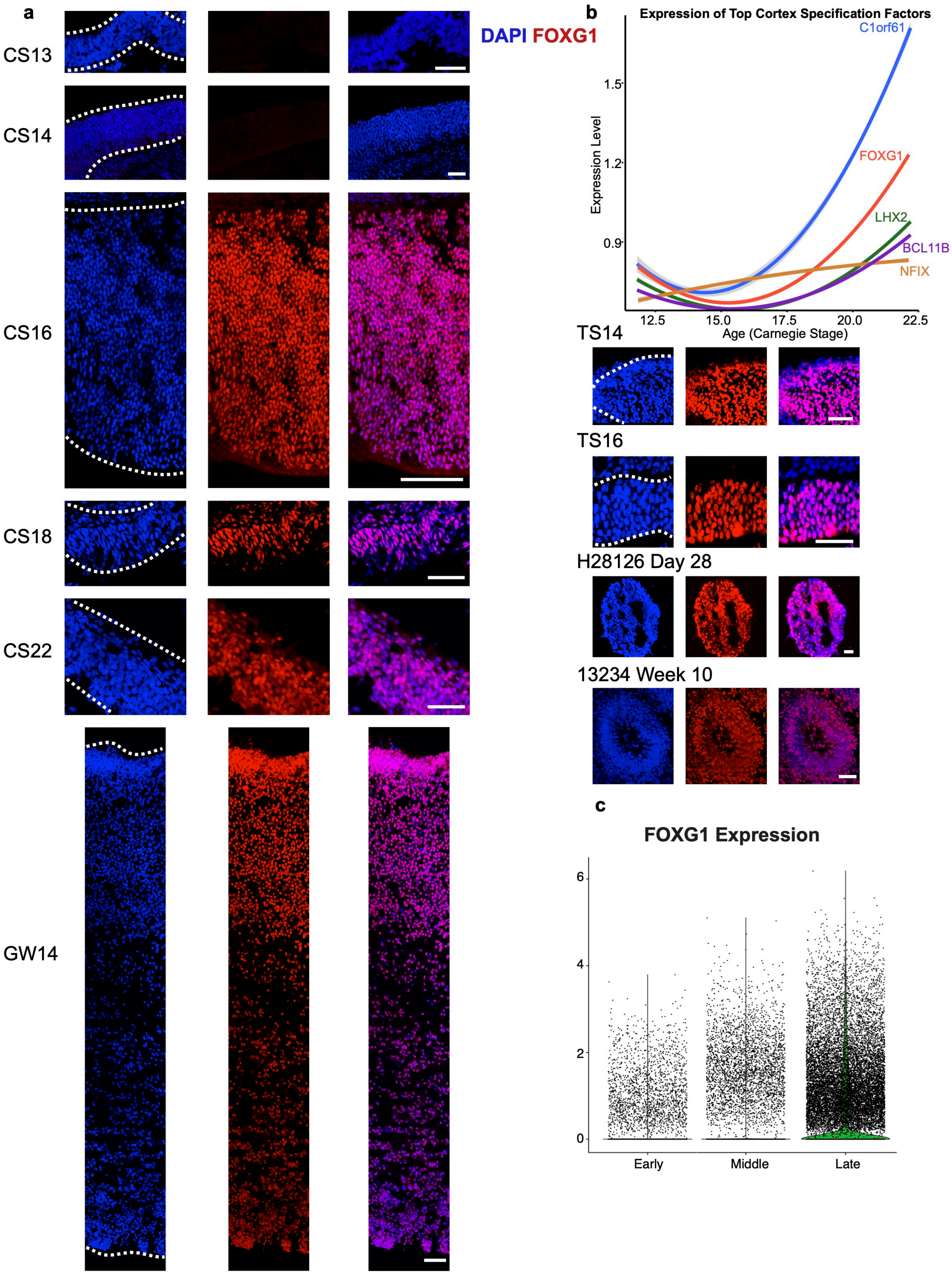

a
CS13
CS14
CS16
CS18
CS22
GW14
DAPI FOXG1
b
Expression of Top Cortex Specification Factors
C1orf61
FOXG1
LHX2
BCL11B
NFIX
Expression Level
1.5
1.2
0.9
12.5
15.0
17.5
20.0
22.5
Age (Carnegie Stage)
TS14
TS16
H28126 Day 28
13234 Week 10
c
FOXG1 Expression
6
4
2
0
Early
Middle
Late

**Supplementary Figure 11. FOXG1 Expression in Developing Human Cortex a)** Immunostaining for DAPI (nuclei in blue) and FOXG1 (red) showing that FOXG1 is not expressed at CS13 or CS14, but is expressed in the adjacent sections used for CS16, CS18, CS22, and GW14 immunostaining validations in this study. Corresponding mouse (TS14, TS16) and cerebral organoid (H28126 Day 28 and 13234 Week 10) samples demonstrate FOXG1 expression also. Each sample was immunostained one time. All scale bars are 50 $\mu$M. **b)** Differentially expressed genes from CS22 were used to identify the most specific cortical genes, and they were plotted across age using ggplot smoothing method “lm”. This plot uses a regression to determine the mean center of the line with the smallest residual. Gray region surrounding each bar shows the 95% error confidence for this linear regression (n = 58,145 cells from 10 individuals).

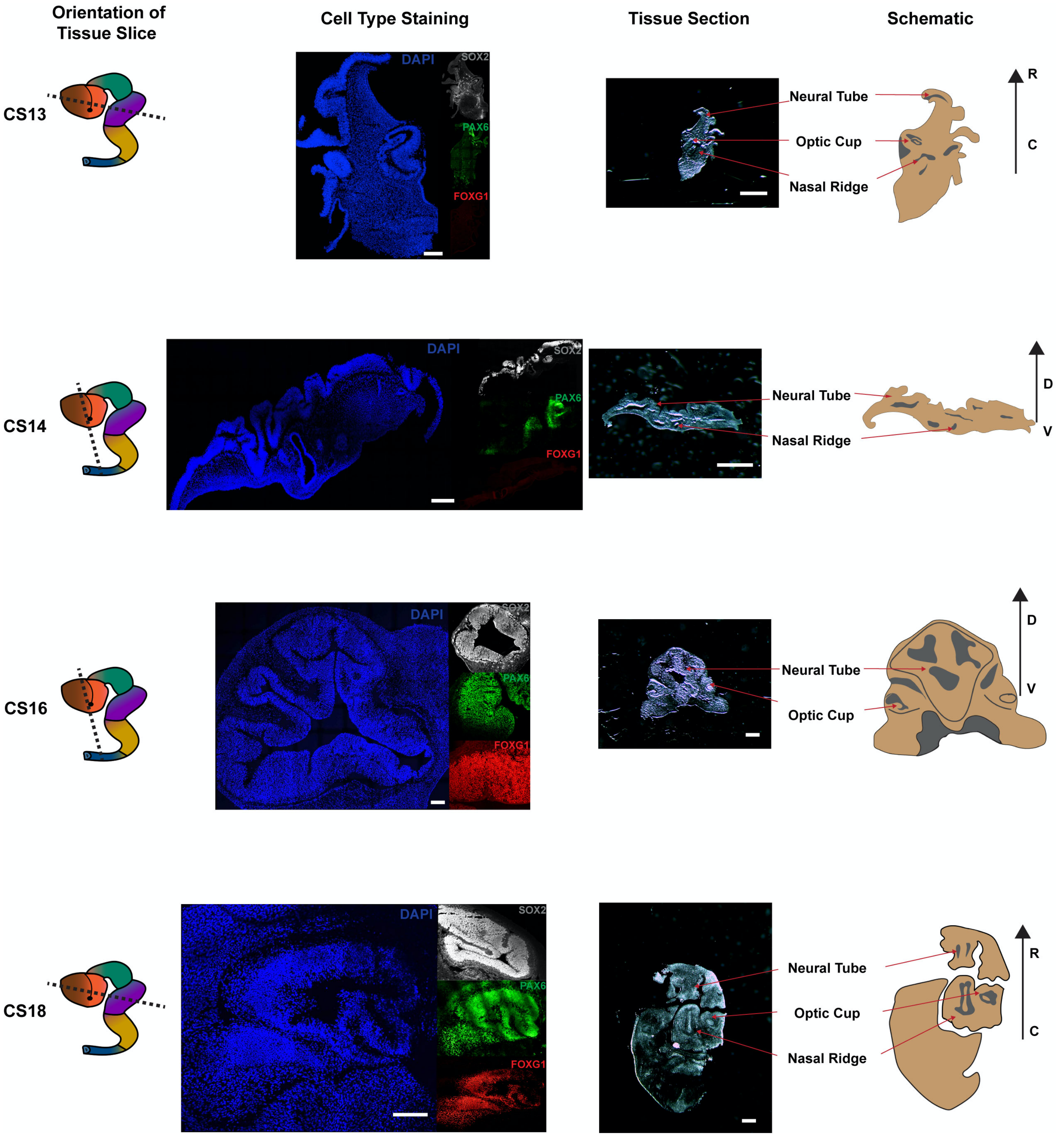


**Supplementary Figure 12. Anatomy of the Developing Human Brain** Samples were sectioned along the orientation of tissue slice (dotted black line) for CS13, CS14, CS16 and CS18. Unstained brightfield images of tissue sections reveal the developing neural tube, optic cup and nasal ridge (red arrows) located caudal to the sections used for imaging in all figure panels. Cell type staining of nuclei (DAPI, blue), SOX2-positive cells (gray), PAX6-positive cells (green) and FOXG1-positive cells (red), reveal similar anatomical structures. Schematic of the tissue sections describe anatomical position of slice: rostral (R), caudal (C), dorsal (D), ventral (V). Each sample was immunostained one time. All cell type staining scale bars are 200 μm. All tissue section image scale bars are 800 μm.

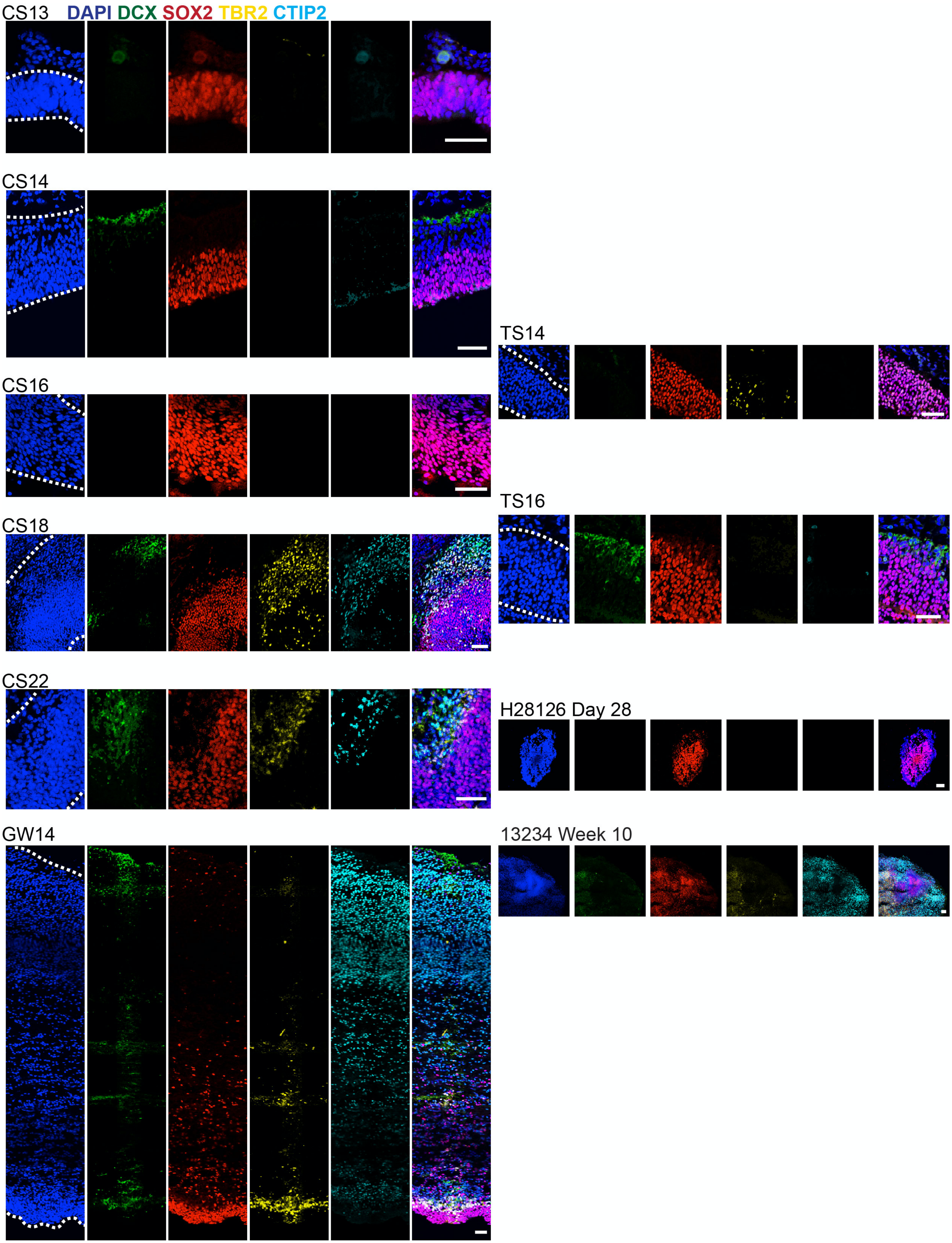


**Supplementary Figure 13. Major Progenitor and Neuronal Populations in the Developing Human Brain**

Immunostaining of progenitors (SOX2, red), IPCs (TBR2, yellow), newborn neurons (DCX, green), maturing neurons (CTIP2, cyan). Nuclei are labeled with DAPI in blue. Progenitors are ubiquitous at CS13, 14, 16, 18, 22 and widely prevalent in the ventricular zone at GW14. Neurogenesis appears to begin as early as CS14, and maturing neurons are observable at CS16. IPCs, which are indicative of indirect neurogenesis, do not emerge until CS18. Mouse samples indicate that neurogenesis begins as early as TS16 as indicated by DCX staining with very sparse TBR2 staining at this age. Cerebral organoid sample (H28126) demonstrates prevalent progenitor staining with very little neuronal or IPC staining at Day 28, but by Week 10 (13234) there is abundant TBR2 and CTIP2 staining. Each sample was immunostained one time. All scale bars are 50 μM.

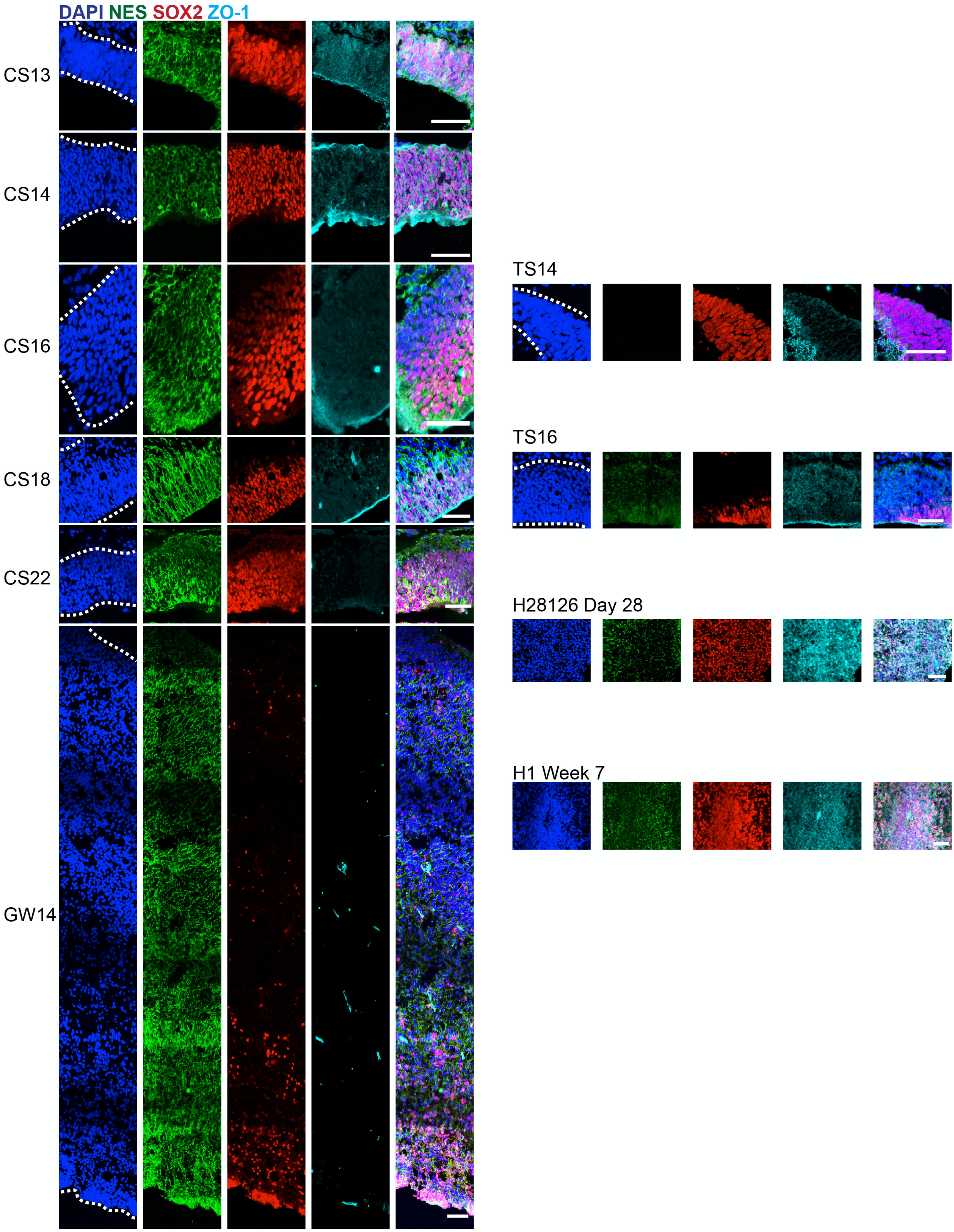


**Supplementary Figure 14. Neuroepithelial and Radial Glia Populations in the Developing Human Brain**

Immunostaining for progenitor populations (SOX2, red) including neuroepithelial cells (ZO-1, cyan) and radial glia (NES, green). Nuclei are labeled with DAPI in blue. From CS13 to CS16 there is a gradual transition from primarily ZO-1 positive to almost entirely NES+ SOX2+ cells. By GW14, ZO-1 only marks presumed vasculature. Mouse samples demonstrate prominent staining of NES by TS16. ZO-1 staining is localized to the ventricular edge in both TS14 and TS16 samples. Cerebral organoid samples (H28126 and H1) shows prominent and diffuse NES and ZO-1 staining at Day 28 and Week 7. Each sample was immunostained one time. All scale bars are 50 μM.

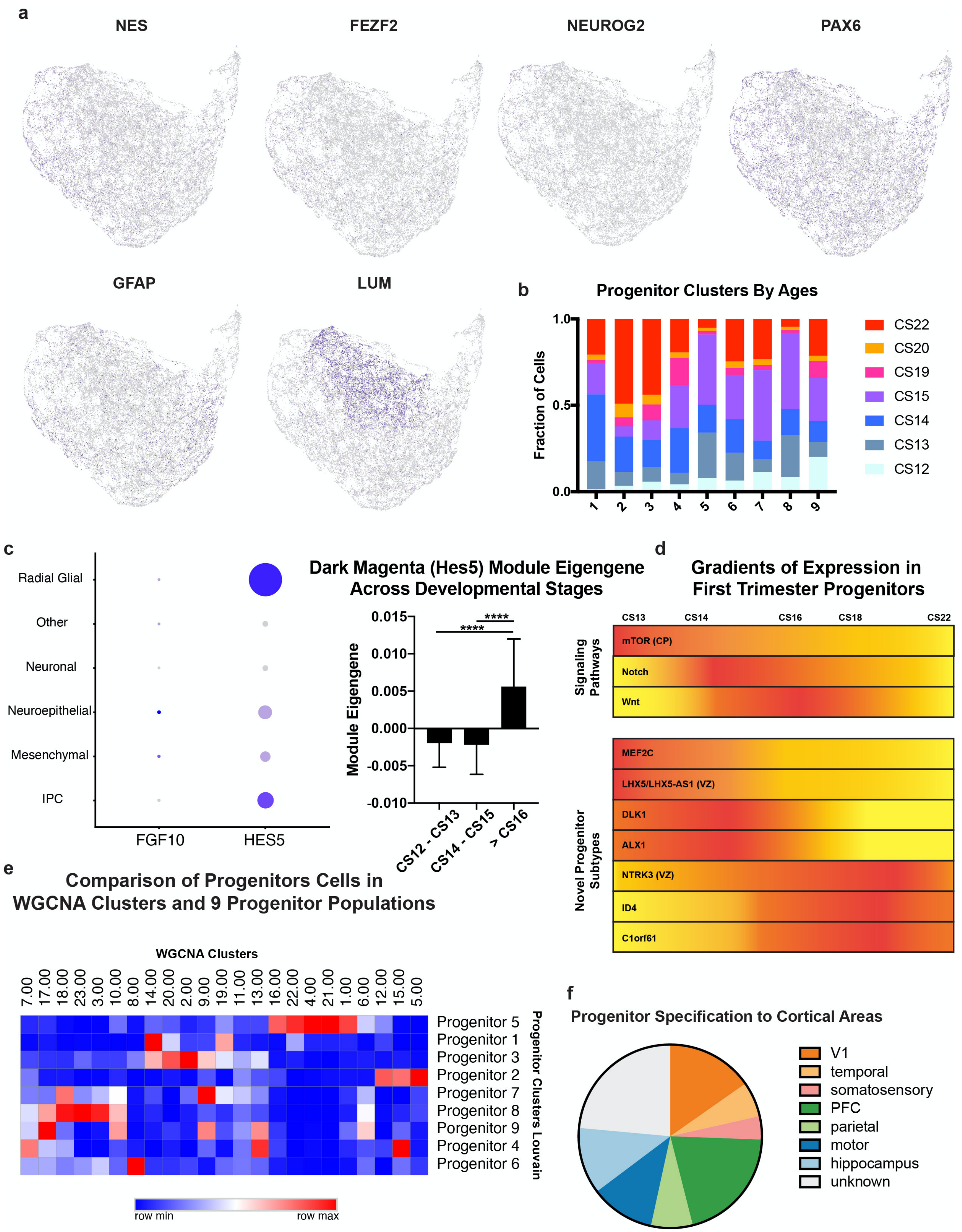


**Supplementary Figure 15. Additional Characterization of Progenitor Clusters a)** Feature plots of genes showing transition to radial glia (*NES*) which is also characterized by expression of neurogenic genes (*FEZF2, NEUROG2*). Most of the progenitors express *PAX6*, and some express *GFAP* which is not expected to be expressed at these early timepoints. *LUM* marks the mesenchymal progenitors. **b)** Stacked bar chart shows the distribution of the ages in each of the progenitor clusters. Progenitor clusters 5 (LUM cluster) and 8 (DLK1 cluster) are almost entirely from early samples. **c)** Dotplot on the left shows *FGF10* spike in neuroepithelial cells and *HES5* enrichment in radial glia. Box and whisker plot (min to max) shows the distribution of module eigengenes for the dark magenta module which includes *HES5* and related genes. One sided t-test was used without multiple hypothesis testing to test the statistical difference between cells older than CS16 (n = 8294 cells from 4 individuals) and the CS14-15 (n = 17,056 cells from 4 individuals) (p-value = $5.02e^{-15}$) or the CS12-13 (n = 7,667 cells from 4 individuals) (p-value = $5.17e^{-29}$). **d)** Patterns of expression of the signaling pathways and novel progenitor cell types. mTOR signaling in the cortical plate (CP) diminishes by the end of the first trimester but reappears in the second trimester to label oRGs. Both *LHX5/LHX5-AS1* and *NTRK3* demonstrate different patterns of expression across the ventricular zone (VZ) and CP. *LHX5/LHX5-AS1* diminishes in the VZ but persists in the CP; while NTRK3 expression peaks at CS18 in the VZ then diminishes, but persists in the CP. **e)** Clustering with WGCNA networks results in 23 clusters that correspond strongly to the nine progenitor subtypes highlighted in this study. **f)** Characterization of progenitor identity in terms of presumptive cortical area shows that most progenitors express genes that correspond to excitatory neuron cortical area during second trimester.

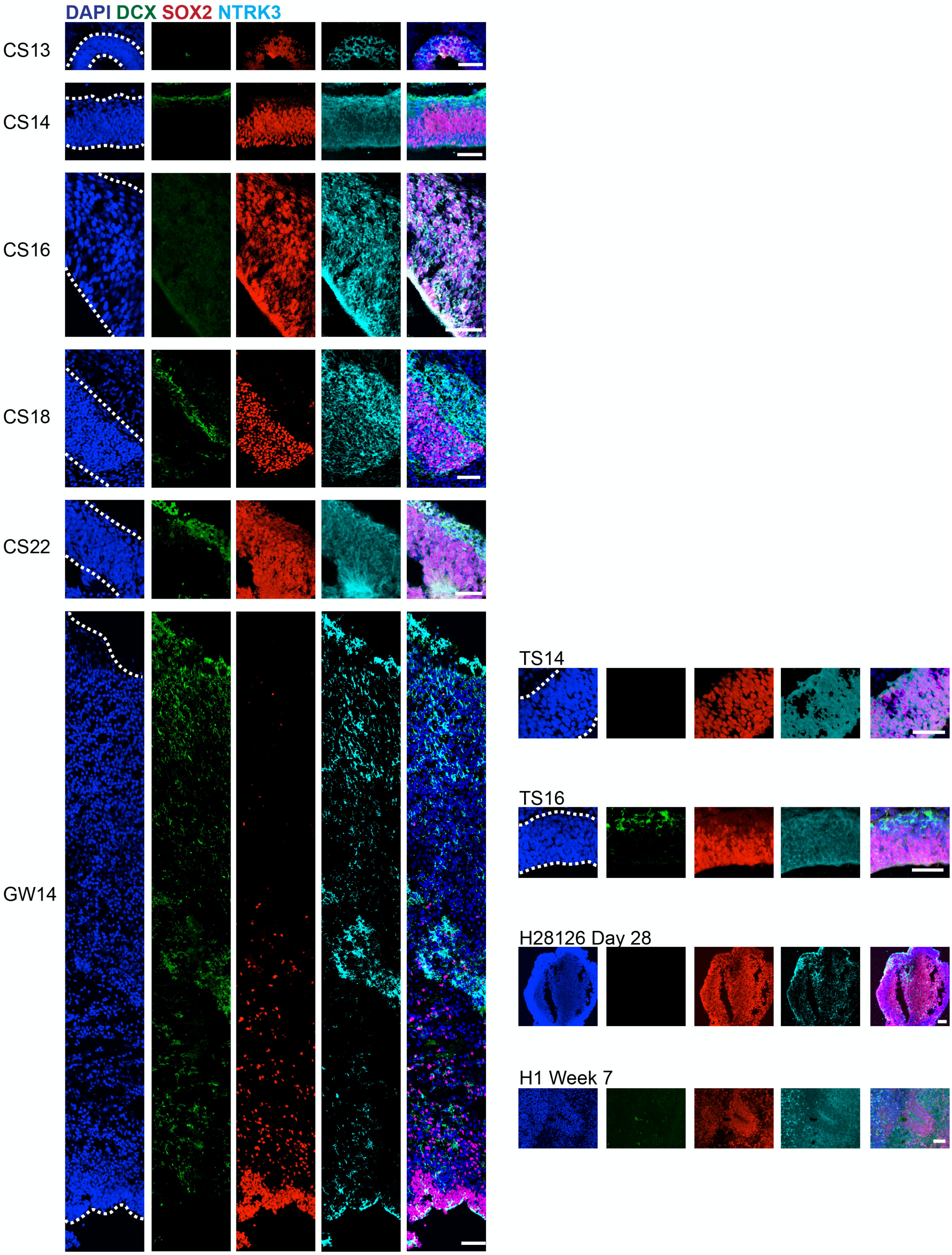

DAPI DCX SOX2 NTRK3
CS13
CS14
CS16
CS18
CS22
GW14
TS14
TS16
H28126 Day 28
H1 Week 7

**Supplementary Figure 16. NTRK3 Expression in the Developing Human Brain** NTRK3 is a unique marker of Progenitor Cluster 3. Immunostaining of NTRK3 (cyan) shows early co-localization with progenitors (SOX2) but at later timepoints the expression pattern more closely coincides with neurons (DCX, green) as has been described in the literature. Nuclei are marked by DAPI in blue. NTRK3 is most widely expressed at CS13, CS14, CS16, CS18, and CS22. At GW14, its expression is restricted to subsets of neurons. Mouse samples illustrate diffuse NTRK3 staining at TS14 and TS16, colocalizing with SOX2+ cells. At Day 28 (H28126), the cerebral organoid sample has diffuse NTRK3 staining with SOX2 colocalization, which persists until Week 7 (H1). Each sample was immunostained one time. All scale bars are 50 μM.

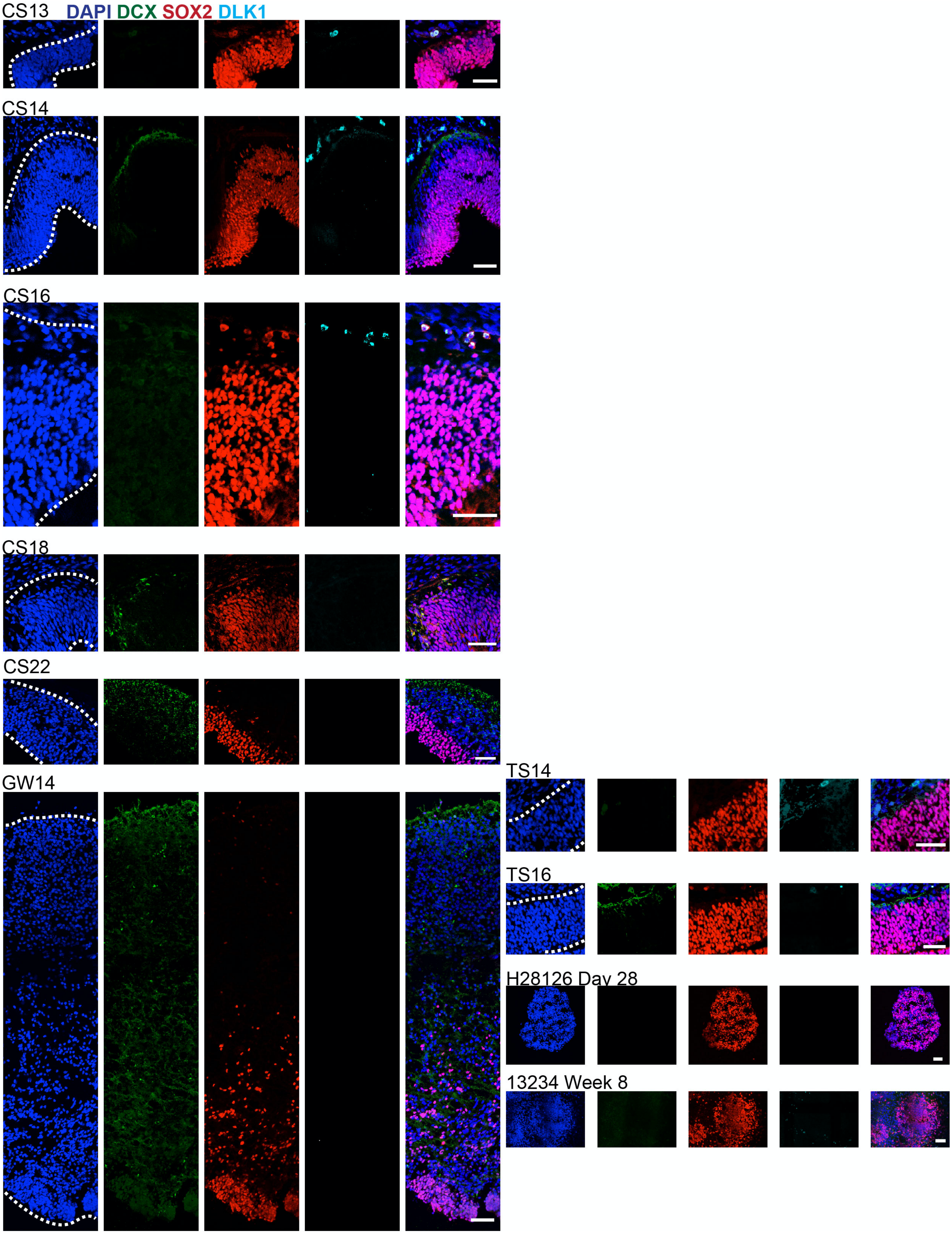
CS13 DAPI DCX SOX2 DLK1
CS14
CS16
CS18
CS22
GW14
TS14
TS16
H28126 Day 28
13234 Week 8

**Supplementary Figure 17. DLK1 Expression in the Developing Human Brain** DLK1 is a marker of early progenitor clusters. Immunostaining for DKL1 (cyan) is sparse but present at CS13, CS14, and CS16 but disappears by CS18 and is not visible in CS22 or GW14. DLK1 co-localizes with progenitors expressing low SOX2 (red), but not neurons (DCX, green). Immunostaining for DLK1 is also present in the TS14 and TS16 mouse samples with no SOX2 colocalization. Cerebral organoid sample (H28126) shows no immunostaining of DLK1 at Day 28 (H28126) or Week 8 (13234). Nuclei are marked by DAPI in blue. Each sample was immunostained one time. All scale bars are 50 μM.

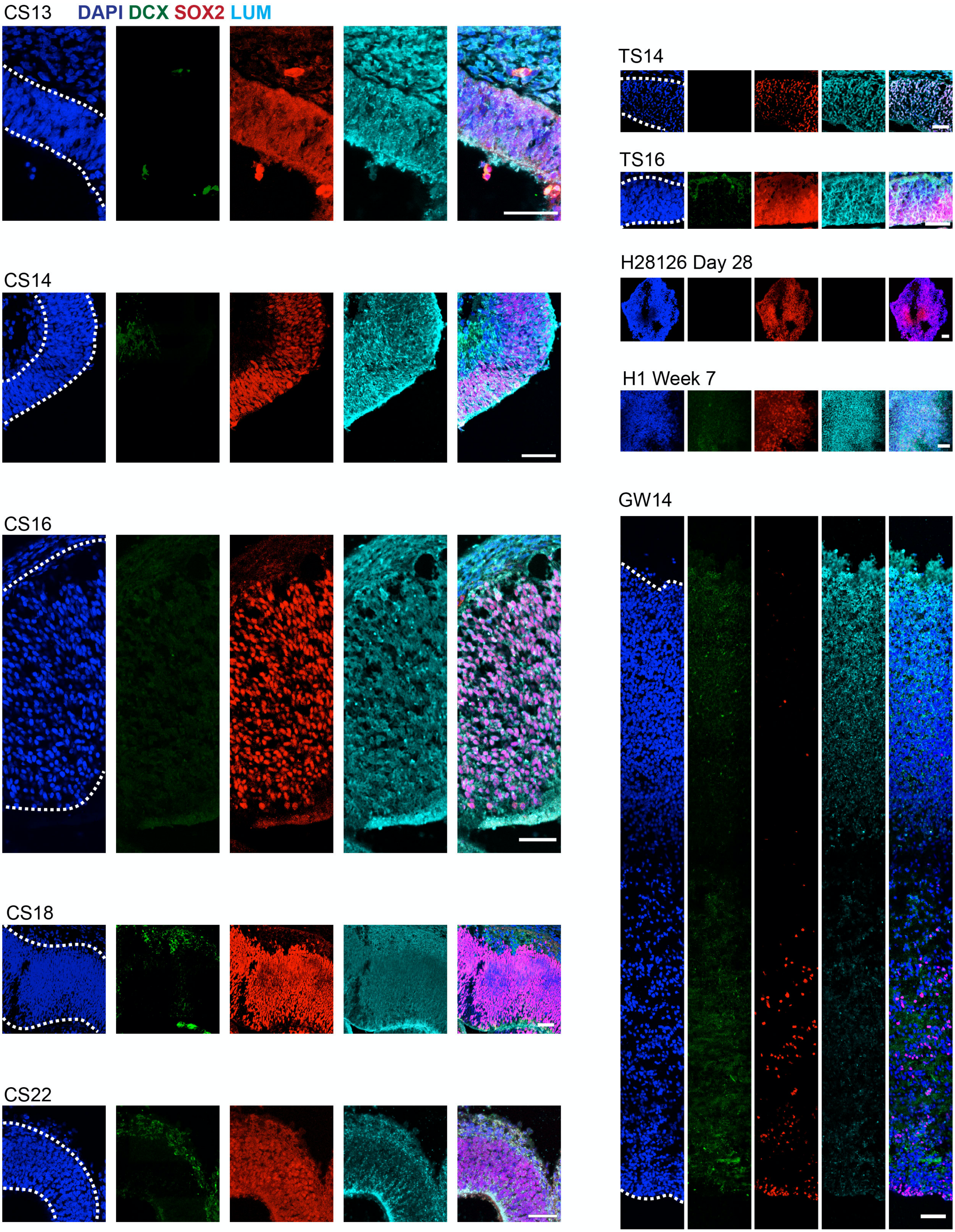
CS13
DAPI DCX SOX2 LUM
CS14
CS16
CS18
CS22
TS14
TS16
H28126 Day 28
H1 Week 7
GW14

**Supplementary Figure 18. LUM Expression in the Developing Human Brain** LUM is a marker for the mesenchymal cell population in the early human cortex, and also a marker for Progenitor Cluster 5. Lumican (LUM, cyan) is a secreted protein and is ubiquitous across all time points. As it is secreted it does not co-localize with either progenitors (SOX2, red) or neurons (DCX, green) but appears to be between cells. There is also prevalent LUM staining in the mouse samples at TS14 and TS16, but there is no staining for LUM in the cerebral organoid sample (H28126) at Day 28. However, by Week 7 (H1) there is diffuse LUM staining. Nuclei are marked by DAPI in blue. Each sample was immunostained one time. All scale bars are 50 μM.

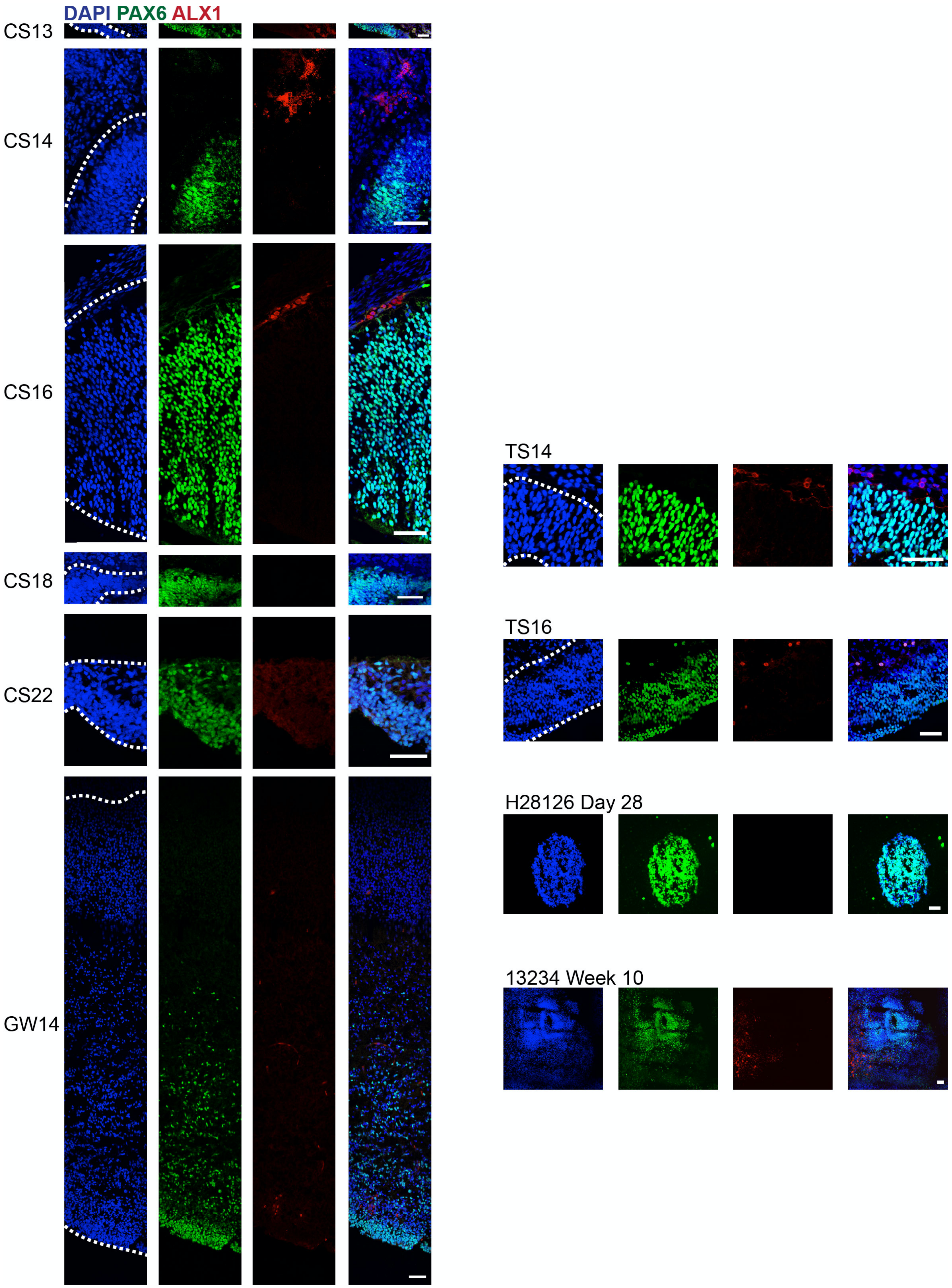

DAPI PAX6 ALX1
CS13
CS14
CS16
CS18
CS22
GW14
TS14
TS16
H28126 Day 28
13234 Week 10

**Supplementary Figure 19. ALX1 Expression in the Developing Human Brain** ALX1 is a marker for the mesenchymal cell population in the early human cortex, and also a marker for Progenitor Cluster 5. ALX1 (red) is sparsely expressed at near the cortical edge at CS13, 14, and 16. Some of the cells also express cortical progenitor markers such as PAX6 (green). ALX1 is no longer present in the cortex at CS18 or 22, but can be found in certain extracortical cells. At GW14 ALX1 marks only presumed vasculature. TS14 and TS16 mouse samples demonstrate extracortical ALX1 staining. There was no staining for ALX1 in the cerebral organoid (H28126) sample at Day 28. However, by Week 10 (13234) there is abundant staining for ALX1. Nuclei are marked by DAPI in blue. Each sample was immunostained one time. All scale bars are 50 μM.

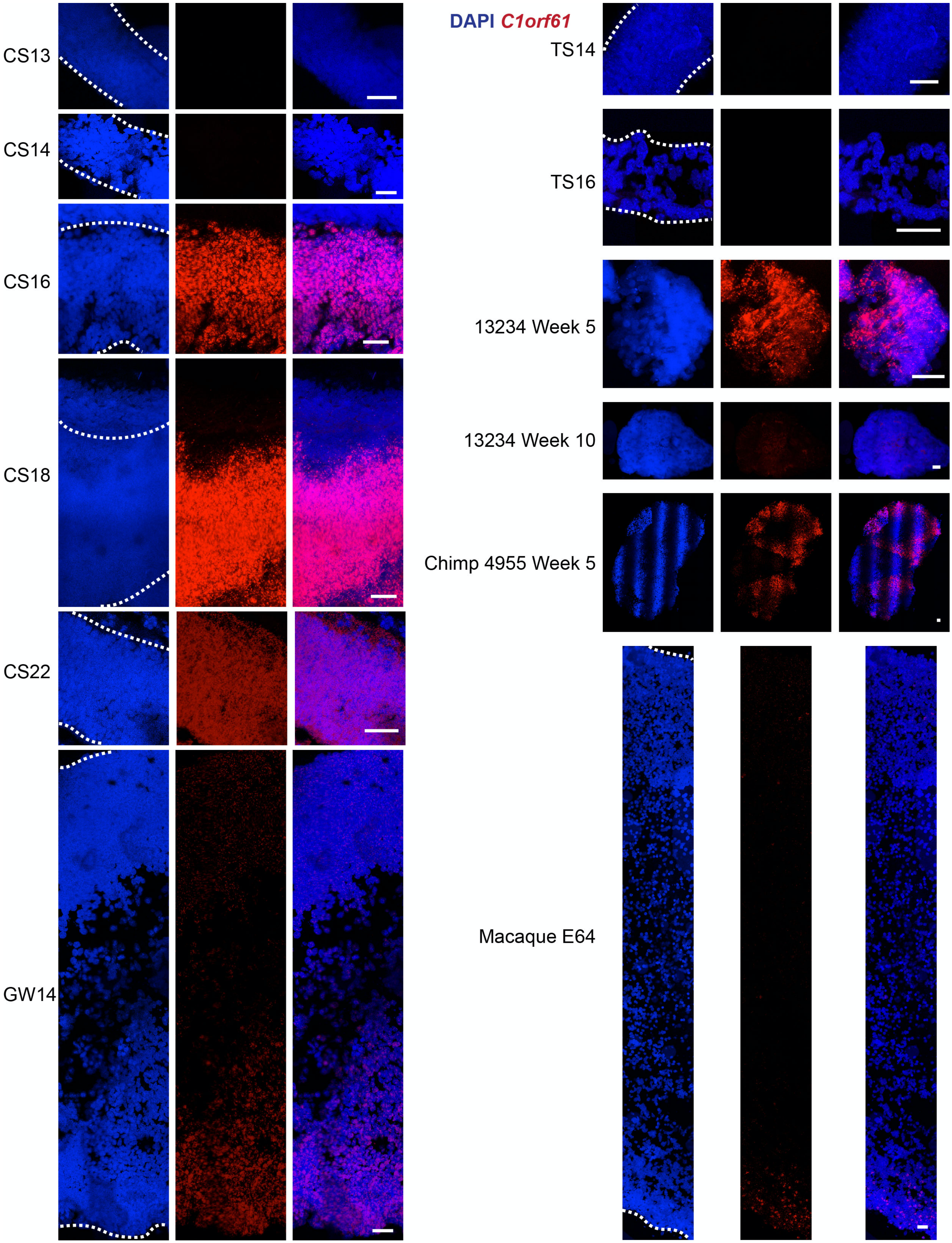


**Supplementary Figure 20. C1orf61 RNA Expression in the Developing Human Brain** *C1orf61* RNA expression minics the same pattern of expression as the protein (SFig 15). *C1orf61* (red) RNA is expressed in a large proportion of progenitors at CS16, 18 and 22 using fluorescent *in situ* hybridization. By GW14 its expression is much lower and also marks a subset of neurons. This is somewhat consistent across primates, as the E64 macaque demonstrates much lower *C1orf61* expression. There is no *C1orf61* expression in the mouse samples at either TS14 or TS16. However, there is diffuse expression for *C1orf61* in the cerebral organoid sample (13234) at Week 5 that becomes less abundant by Week 10 (13234). At Week 5, the chimp organoid (4955) demonstrates diffuse *C1orf61* expression. Nuclei are marked by DAPI in blue. Each sample was processed and stained one time. All scale bars are 50 μM.

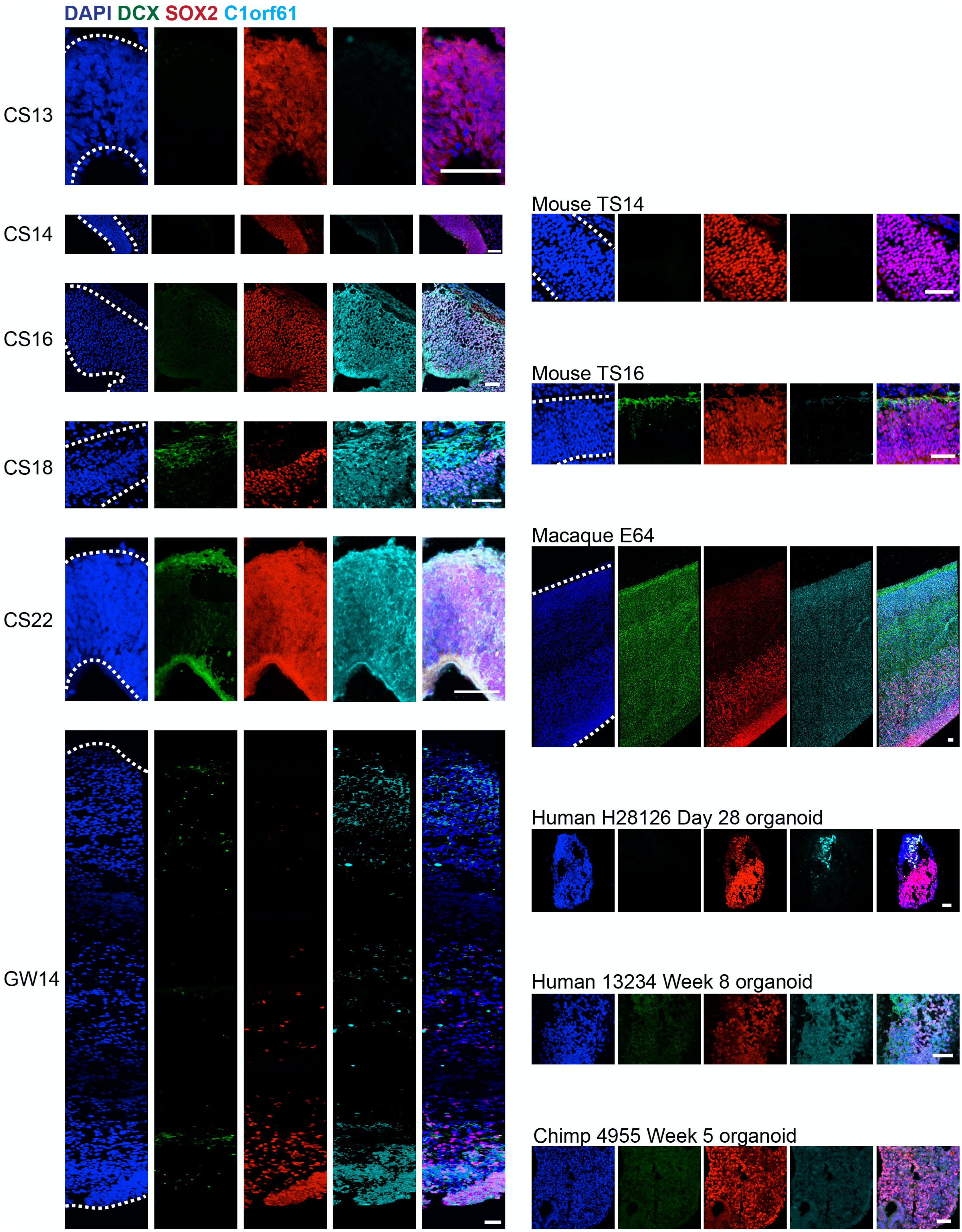


**Supplementary Figure 21. C1orf61 Expression in the Developing Human Brain** C1orf61 is a marker of Progenitor Cluster 4 and also ramps up significantly over the course of human cortical development (SFig 5b).

It does not appear to be expressed at similar timepoints of mouse development (Fig 4). By immunostaining, C1orf61 (cyan) marks a large proportion of progenitors (SOX2, red) at CS16, 18 and 22. By GW14 its expression is much lower and also marks a subset of neurons (DCX, green). This is consistent across primate species, as the E64 macaque demonstrated diffuse C1orf61 staining. There was no C1orf61 staining in the mouse samples at either TS14 or TS16. However, there was sparse staining for C1orf61 in the cerebral organoid sample (H28126) at Day 28 that becomes more abundant by Week 8 (13234). At Week 5, the chimp organoid (4955) demonstrates low but diffuse C1orf61 staining. Nuclei are marked by DAPI in blue. Each sample was immunostained one time. All scale bars are 50 μM.

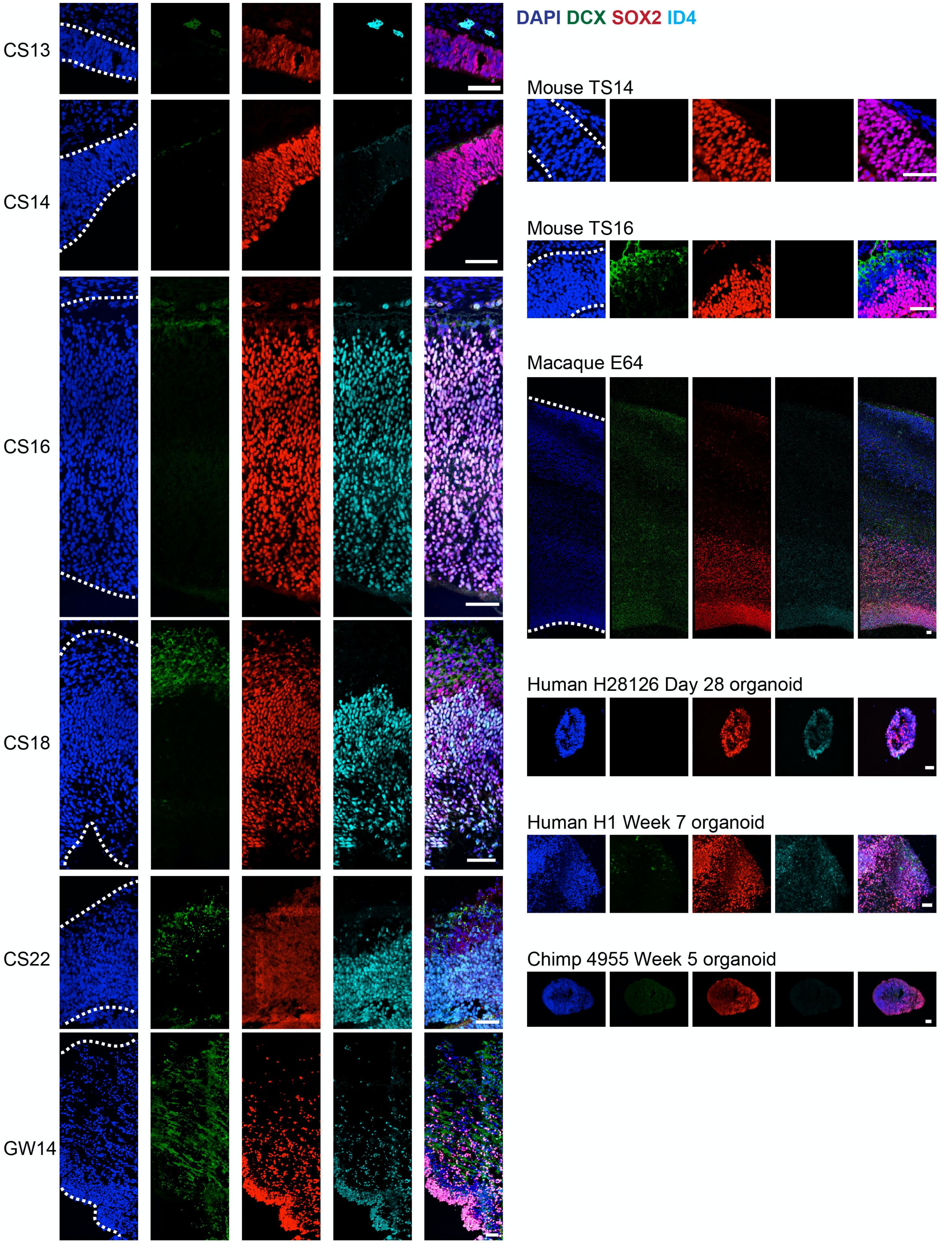
DAPI DCX SOX2 ID4
CS13
CS14
CS16
CS18
CS22
GW14
Mouse TS14
Mouse TS16
Macaque E64
Human H28126 Day 28 organoid
Human H1 Week 7 organoid
Chimp 4955 Week 5 organoid

**Supplementary Figure 22. ID4 Expression in the Developing Human Brain** ID4 is a marker of Progenitor Cluster 7. It does not appear to be expressed at similar timepoints of mouse development (Fig 4). By immunostaining, ID4 (cyan) marks a large proportion of progenitors (SOX2, red) at CS16, 18 and 22 and GW14. This pattern of staining is consistent in the E64 macaque sample. There is no ID4 staining in the mouse samples at either TS14 or TS16. However, there is staining for ID4 in the cerebral organoid sample (H28126) at Day 28, and that becomes more pronounced by Week 7 (H1). At Week 5, the chimp organoid (4955) demonstrates no ID4 staining. Nuclei are marked by DAPI in blue. Each sample was immunostained one time. All scale bars are 50 $\mu$M.

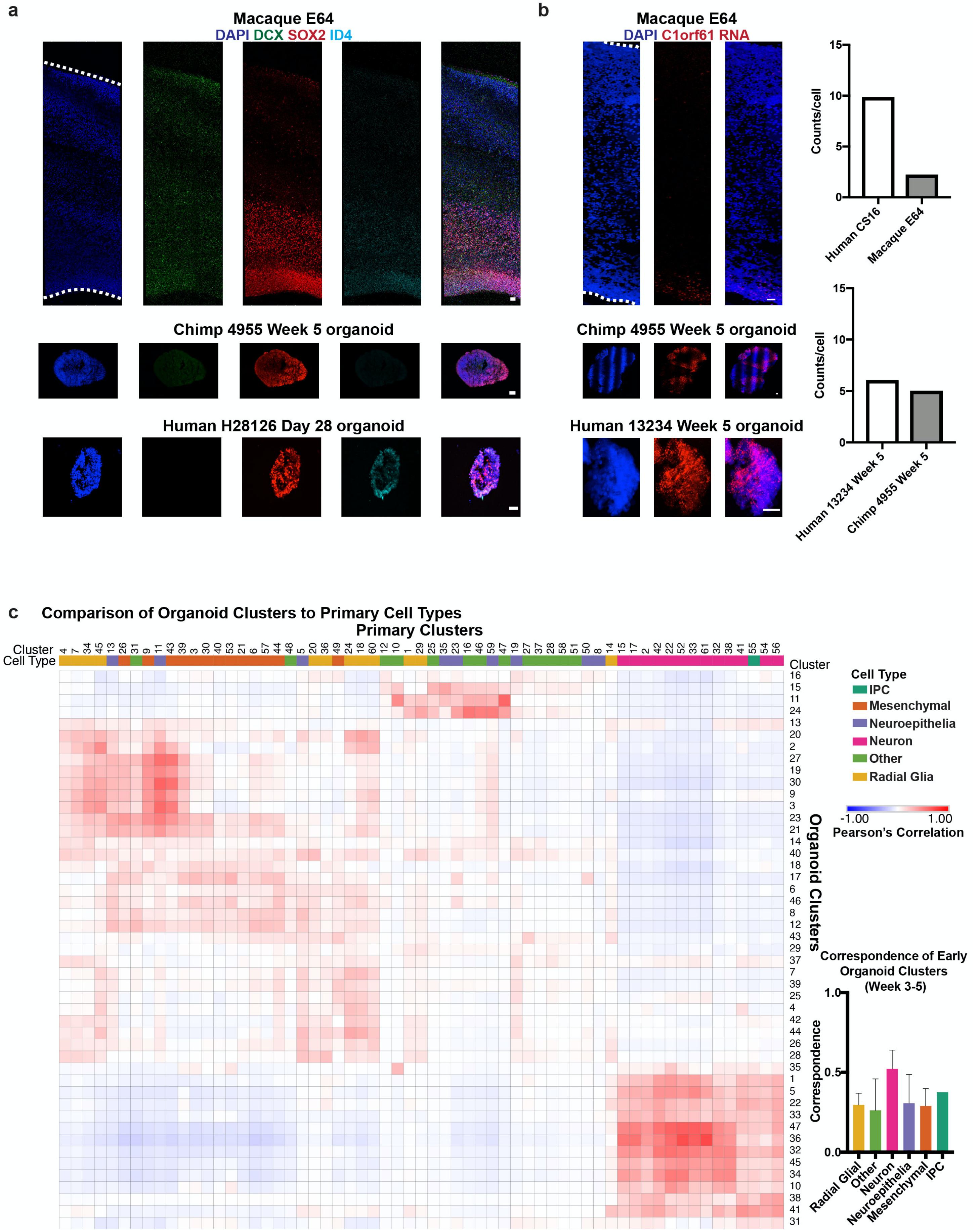


**Supplementary Figure 23. Non-human Primate and Organoids Compared to the Primary Developing Human Cortex a)** ID4 expression is present but faint in the E64 macaque sample. There appears to be no ID4

staining by Week 5 in the chimp organoid (4955). However, there is abundant staining of ID4 in the human organoid sample (H28126) by Day 28. **b)** *C1orf61* RNA expression is also present but faint in the E64 macaque sample, as compared to the CS16 human, which demonstrates higher expression. There appears to be diffuse and abundant RNA expression, however, of *C1orf61* in both the chimp (4955) (n = 538 cells from 1 individual) and human (13234) (n = 707 cells from 1 individual) organoid samples by Week 5. Each sample was processed and stained one time. All scale bars are 50 μm. **c)** Heatmap showing correlation between organoid clusters from week 3 and 5 organoids (from Bhaduri et al, 2020) to primary cells from this study. Correlations are performed using Pearson correlations between cluster marker sets, and summary metrics across all primary clusters in each cell type category are shown in the bar plot (mean with standard deviation). Excitatory neurons are the most highly preserved cell type in organoids, but progenitors, including neuroepithelial cells and radial glia, are substantially different between our primary data and organoid datasets.

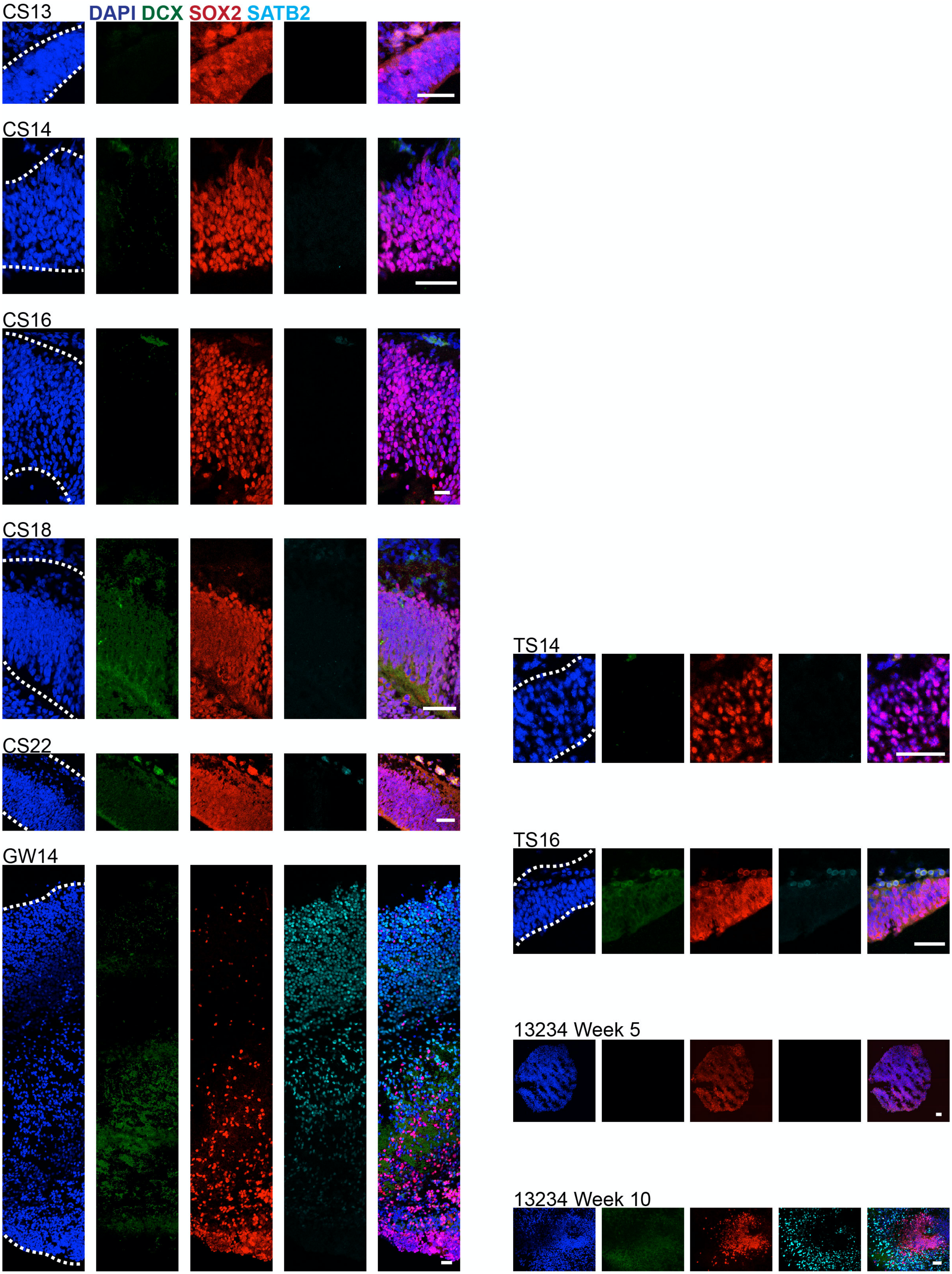
CS13
DAPI DCX SOX2 SATB2
CS14
CS16
CS18
CS22
GW14
TS14
TS16
13234 Week 5
13234 Week 10

**Supplementary Figure 24. SATB2 Expression in the Developing Human Brain** SATB2 is a marker for upper layer neurons and is highly enriched in a subset of the neuronal population of this dataset. SATB2 (cyan) only marks subpopulations of neurons (DCX, green) by GW14. SATB2 staining was not present any earlier. Immunostaining for SATB2 demonstrated very faint staining in TS16 mouse sample. Cerebral organoid samples (H28126 and 13234) show diffuse immunostaining for SATB2 only by Week 10. Progenitors are marked by SOX2 in red. Nuclei are marked by DAPI in blue. Each sample was immunostained one time. All scale bars are 50 μM.

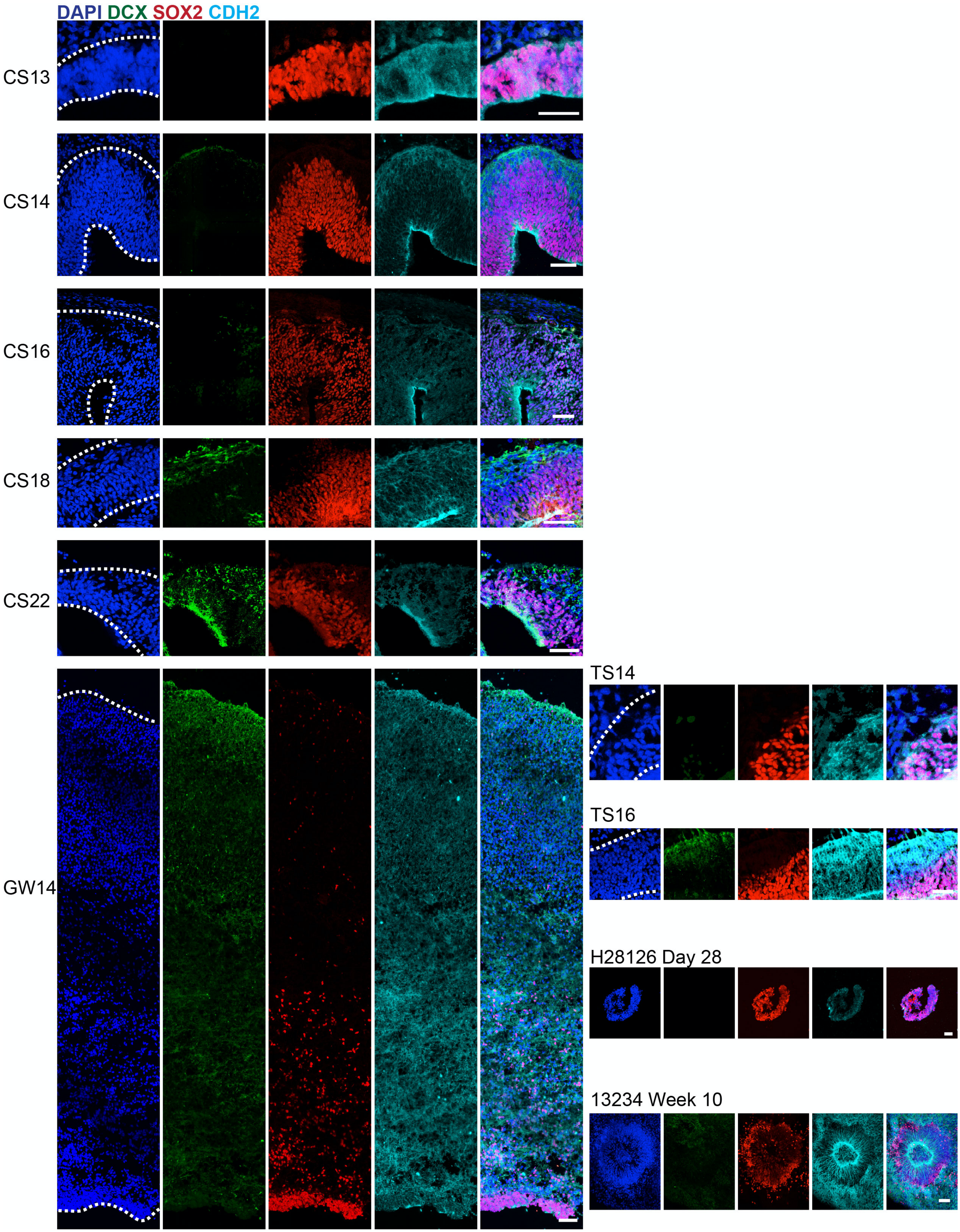


**Supplementary Figure 25. CDH2 Expression in the Developing Human Brain** CDH2 is a marker of Progenitor Cluster 9. CDH2 (cyan) is a cadherin that marks subpopulations of both progenitors (SOX2) and some neurons (DCX, green) across the ages sampled. Immunostaining for CDH2 in both TS14 and TS16 mouse samples demonstrate staining throughout the cortical span. Cerebral organoid samples (H28126 and 13234) also shows diffuse immunostaining for CDH2 at Day 28 and Week 10. Nuclei are marked by DAPI in blue. Each sample was immunostained one time.  All scale bars are 50 μM.

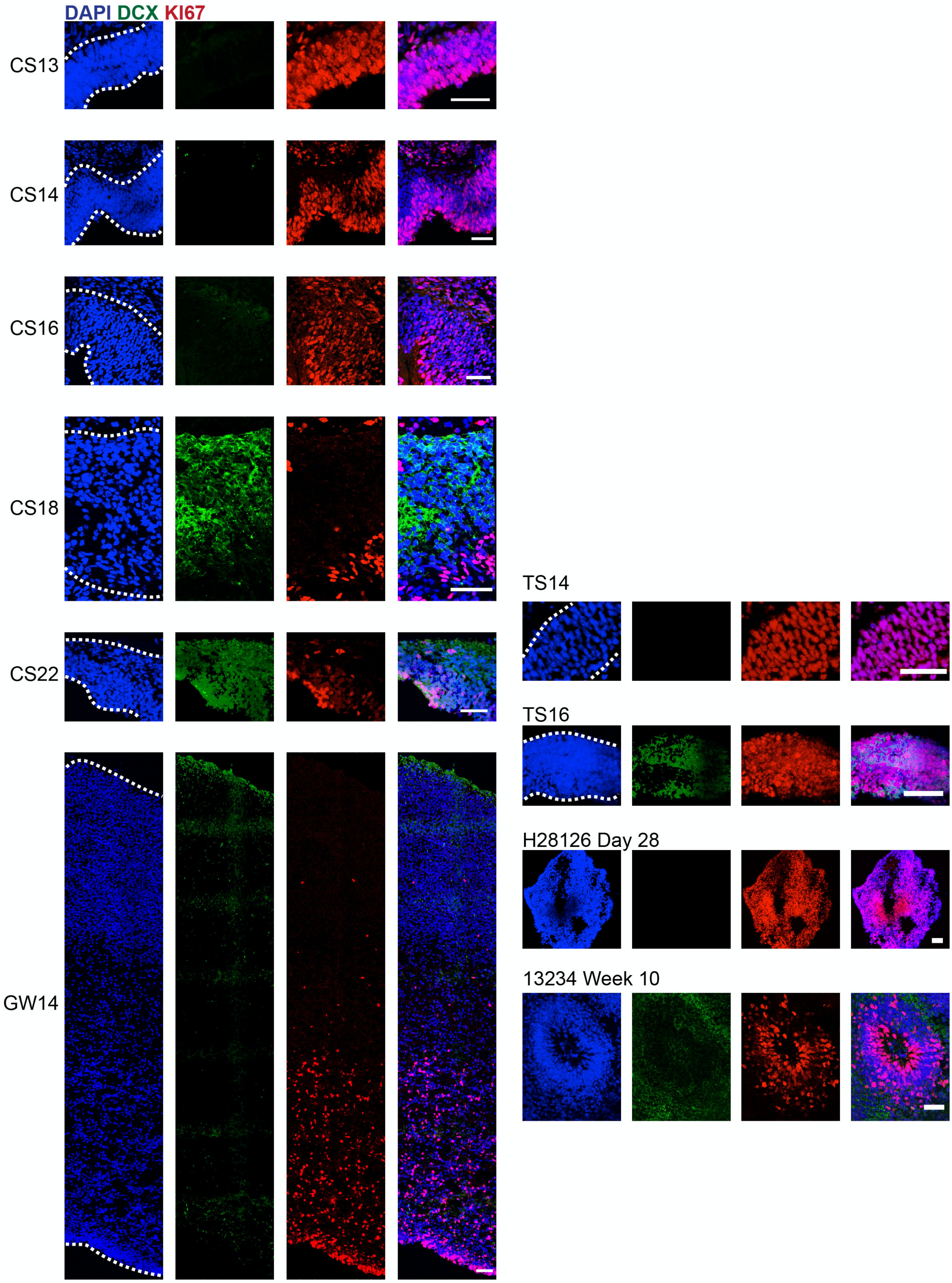
DAPI DCX KI67
CS13
CS14
CS16
CS18
CS22
GW14
TS14
TS16
H28126 Day 28
13234 Week 10

**Supplementary Figure 26. Dividing Progenitor Cells During Early Stages of Human Cortical Development** Immunostaining for KI67 (red) marks dividing progenitors. These cells are nearly ubiquitous in CS13, 14, and 16, and mark a large percentage of the progenitor zone (but not neurons, DCX, green) in CS18 and 22. GW14 has a lower number of dividing cells, but they are still very visible in the ventricular zone. Mouse samples demonstrate ubiquitous KI67 staining at both TS14 and TS16. Cerebral organoid samples (H28126 and 13234) also shows broad KI67 staining at Day 28 up until rosette formation at Week 10 (13234). Nuclei are marked by DAPI in blue. Each sample was immunostained one time. All scale bars are 50 μM.
